# Neuronal p38a knockout protects against neurological consequences following repetitive mild traumatic brain injury

**DOI:** 10.64898/2026.02.26.708089

**Authors:** Chenxing Li, Sydney E. Triplett, Martin N. Griffin, Abigail L. Holberton, Adnan Kadragic, Felix G. Rivera Moctezuma, Sia J. Saheba, Paul F. Saah, Paul I. Sanz, Janie C. Lee, Rohan Wadhwani, Drew Dawson, Sophie E. Lunt, Maya Chigurupati, Erin M. Buckley, Levi B. Wood

## Abstract

Mild traumatic brain injuries (mTBI) can substantially impact quality of life, and repetitive mTBIs (rmTBI) can amplify injury effects compared to a single injury. However, effective clinical treatments remain elusive, largely due to an incomplete understanding of the underlying injury mechanisms. Neuroinflammation has emerged as a key contributor to worse functional outcomes after mTBI/rmTBI. While microglia are traditionally viewed as primary mediators of post-injury inflammation, accumulating evidence suggests neurons play an immunomodulatory role in initiating the rmTBI inflammatory cascade through activation of intracellular proinflammatory pathways like p38 MAPK and secretion of cytokines that, in turn, stimulate microglial activation. Here, we tested whether inducible neuronal p38α knockout protects against functional, immune, and cerebrovascular consequences of a weight-drop closed head injury model of rmTBI. A battery of functional assays was conducted 4 weeks post-injury, and tissues were collected at both 4 hours and 4 weeks following final CHI. In males, neuronal p38α knockout protected against injury-induced depressive-like behavior, hyperactivity, synaptic loss, microglial reactivity, cytokine upregulation, and reduction in cerebral blood flow. In females, neuronal p38α knockout protected against risk-taking behavior and partially protected against cytokine upregulation but had limited effect on microglial reactivity and cerebral blood flow. Together, these findings identify neuronal p38α as a sex-dependent driver of rmTBI-associated neurological consequences, and they support neuronal p38α-immune signaling as a mechanistically relevant therapeutic target for future studies.

## Introduction

Every year, approximately 3 million people suffer from a mild traumatic brain injury (mTBI) in the United States alone, resulting in a healthcare burden of approximately $17 million dollars^1,2^. Although mTBIs are not life threatening, the neurological consequences can persist for months or even years post-injury^3–6^. Moreover, repetitive mild TBIs (rmTBI)s, especially when sustained within a vulnerable recovery window, can amplify deficits compared to a single injury^7–9^. For example, recent work found that individuals with four or more mTBIs exhibited significant long-term deficits in attention, processing speed, and working memory, with progressive decline following each additional injury^8^. Despite the prevalence and impact on quality of life, effective clinical treatments for mTBI/rmTBI remain elusive, largely due to an incomplete understanding of the underlying injury mechanisms.

Over the past decade, neuroinflammation has emerged as a key contributor to worse functional outcomes following mTBI/rmTBI^10–16^. The neuroinflammatory response involves rapid intracellular phospho-protein signaling, cytokine and chemokine release, and glial cell activation. Additional secondary injury processes, including axonal injury, altered neurotransmission, blood–brain barrier disruption, ionic imbalance, and excitotoxicity, further exacerbate the neuroinflammatory effects of injury^17–20^. While microglia have long been recognized as key players in secreting and responding to inflammatory signals^11,21,22^, recent evidence suggests that neurons also contribute to the inflammatory response through activation of intracellular proinflammatory pathways and secretion of cytokines that, in turn, stimulate microglial activation^18,23–25^.

Among the intracellular phospho-protein signaling pathways that are activated after injury, p38 mitogen activated protein kinase (MAPK) has emerged as a key driver of functional outcomes across various mild to severe TBI injury models^11,26–31^. The p38 MAPK phospho-signaling proteins represent a non-canonical portion of the MAPK pathway that responds to extracellular stimuli and stress with known roles in inflammation^26,28,32–34^, synaptic dysfunction^27,33,35^, and neuronal apoptosis^34,36^. In rmTBI, pharmacological inhibition of p38ɑ/β mitigated a broad range of rmTBI-induced deficits from transcript alterations to behavioral deficits^31^. While p38 is expressed in diverse cell types (e.g. microglia, neurons, astrocytes), our work in rmTBI has consistently shown co-labelling between the neuronal marker NeuN and phosphorylated p38 (p-p38) as well as downstream injury-induced cytokines, implicating a role of neuronal p38 in the rmTBI-induced inflammatory cascade. Coupled with the growing body of evidence indicating that neurons actively upregulate and secrete immune mediators^23,24,37–39^, these findings support an immunomodulatory role for neurons in initiating the rmTBI inflammatory cascade.

In this work, we investigated the role of neuronal p38α in driving neurological consequences after rmTBI in neuronal p38α knockout transgenic mice. We specifically chose the p38α (MAPK14) isoform because it is recognized as a key inflammatory kinase that is linked to both neurodegenerative disease and synapo-toxicity^37,40,41^. Using tamoxifen-induced knockout prior to injury, we quantified the effects of neuronal p38α knockout on immune response, synaptic function, microglial phenotype, cerebral hemodynamics, and functional outcomes after a repetitive weight-drop closed head injury (CHI) model. We hypothesized that neuronal p38α knockout would alleviate injury-induced pathological and neurological consequences.

## Materials and Methods

### Animals

All protocols were approved by the Emory University Institutional Animal Care and Use Review Office. Male and female mice were housed in the university animal facility on a 12-hour light/dark cycle. Food and water were provided *ad libitum*.

To achieve neuron-specific conditional p38α knockout, we first crossed tamoxifen-inducible STOCK Tg (Thy1-cre/ERT2-EYFP)HGfng/PyngJ1 mice (Jackson Laboratory Strain#: 012708) with STOCK *Mapk14^tm1.2Lex^*/YiwaJ mice (Jackson Laboratory Strain#: 031129) to generate hemizygous SLICK-H Cre-Thy1^+/Cre-Thy1^; p38α^+/fl^ mice. Hemizygous carriers were then crossed with p38α^fl/fl^ to generate SLICK-H Cre-Thy1^+/Cre-Thy1^; p38α^fl/fl^ mice (abbreviated as CRE herein). Floxed p38α^fl/fl^; SLICK-H Cre-Thy1^+/+^ mice lacking Cre were used as experimental wild type controls (abbreviated as WT herein). Both male and female mice aged to 3-5 months were used for the experiments described in the following section.

### Study protocol

All mice were given 5 daily intraperitoneal injections of tamoxifen (75 mg/kg in corn oil) to initiate Cre-mediated recombination in the CRE animals and to control for potential confounding tamoxifen-induced effect in WT animals. Following injections, mice underwent a minimum 2-week tamoxifen washout period prior to experimental intervention. Mice were randomly assigned to one of two groups: five once-daily closed head injuries (5xCHI) or five once-daily sham injuries. As such, we had a total of 4 experimental groups: 5xCHI + CRE, 5xCHI + WT, sham + CRE, sham + WT. Animals were sacrificed by cervical dislocation under 5% isoflurane (1L/min, 100% oxygen) at either a short (4 hours) or long (∼4 weeks) time point after injury/sham injury. Mice that were sacrificed at the long time point underwent a battery of functional assessments 2-4 weeks following the final CHI. Cerebral blood flow was assessed at baseline and at 4 hours after the final injury/sham injury in all mice. All brain tissues were collected after euthanasia and separated by hemisphere. From the left hemisphere, the frontal cortex, somatomotor cortex, and hippocampus were micro-dissected, flash frozen in liquid nitrogen, and stored at −80°C for molecular analyses. Right hemispheres were fixed in 4% paraformaldehyde, paraffin-processed, and embedded for immunohistochemistry.

### Closed Head Injury model

Injury was induced with a previously described weight drop, closed head injury (CHI) model^9,42^. In brief, mice were anesthetized with 3–5% isoflurane (1 L/min, 100% O₂) for ∼2 minutes, at which time they were removed from anesthesia, positioned on a task wipe (Kimwipes®, Kimberly-Clark, Irving, TX) with their head under a 0.96 m guide tube (McMaster-Carr 49035K85), and grasped by the tail. A 54 g weight was dropped down the tube for impact on the dorsal portion of the head, approximately between the coronal and lambdoid sutures, resulting in anterior–posterior rotation of the head about the neck. Following injury, animals were placed on a warm plate and monitored continuously until they regained righting reflex. None of the injured animals had skull fracture or hemorrhage, consistent with previous reports^43^. Sham-injured mice received the same exposure to anesthesia but were not subject to closed-head injury.

### Protein Analysis

To elucidate the effect of neuronal p38ɑ knockout on molecular changes in the brain, following 5xCHI, we accessed 3 classes of proteins related to pathology (post synaptic marker 95; PSD95), glial phenotype (microglial activation marker CD68 and homeostatic marker TMEM119), and cytokine expression (panel of 18 cytokines/chemokines). A total of 74 samples were collected for these analyses at the short 4 hours post-injury time point (females: n=8 5xCHI + CRE, n=9 5xCHI + WT, n=10 sham + CRE, n=9 sham + WT; males: n=9 5xCHI + CRE, n=10 5xCHI + WT, n=10 sham + CRE, n=9 sham + WT). A total of 92 samples were collected at the long 4 week post-injury time point (females: n=14 5xCHI + CRE, n=11 5xCHI + WT, n=12 sham + CRE, n=11 sham + WT; males: n=10 5xCHI + CRE, n=14 5xCHI + WT, n=11 sham + CRE, n=9 sham + WT). Protein analysis was performed on hippocampal tissues from the left hemisphere lysed in 8M Urea buffer followed by Pierce BCA Protein Assay (Thermo Fisher #23,225).

To quantify synaptic changes, we quantified post synaptic marker 95 (PSD95). Measurements were made via an enzyme-linked immunosorbent assay (ELISA). Tissue lysates were thawed on ice and centrifuged at 4°C for 10 min at 15,500g prior to analysis. After protein concentration for each sample was determined, lysates were diluted to 0.007 μg/μL in 100 ul with respective assay diluents from each kit (Lifespan Biosciences LS-F7142-1). Loaded protein amounts were selected to fall within the linear detection range of absorbance versus protein amount. Buffer-only background signals were subtracted.

To quantify glial phenotype changes, we quantified two microglial markers: macrophage and microglia activation marker cluster of differentiation 68 (CD68)^44^ and microglial homeostatic marker transmembrane protein 119 (TMEM119)^45^. Measurements were made via ELISA. Tissue lysates were thawed on ice and centrifuged at 4°C for 10 min at 15,500g prior to analysis. After protein concentration for each sample was determined, lysates were diluted to 0.1 μg/μL in 50 µl with respective assay diluents from each kit (Lifespan Biosciences LS-F11095 and LS-F52734). Loaded protein amounts were selected to fall within the linear detection range of absorbance versus protein amount. Buffer-only background signals were subtracted.

To quantify cytokines, the Milliplex® MAP Mouse Cytokine/Chemokine 32-plex kit was used (Millipore Sigma, MCYTMAG-70K: Eotaxin, GM-CSF, IFN-γ, IL-1β, IL-2, IL-3, IL-4, IL-6, IL-10, IL-12p70, IL-13, IP-10, MIP-1α, MIP-1β, MIP-2, RANTES, TNF-α, VEGF). After protein concentration for each sample was determined, lysates were adjusted in Milliplex® MAP Assay Buffer to 6µg protein per 12.5 µL. Loaded protein amounts were selected to fall within the linear range of bead fluorescence intensity versus protein concentration for detectable analytes. Buffer-only background signals were subtracted.

### Immunohistochemistry

Immunohistochemistry (IHC) was conducted to visualize and quantify TMEM119 and PSD95. Right brain hemispheres were fixed in 4% paraformaldehyde for 20-24 hours, paraffin processed, embedded, cut into 5 μm sagittal sections, and mounted onto glass slides. On day 1, sections were deparaffinized in xylenes, dehydrated in 100% and then 95% ethanol, and rehydrated with deionized water. Antigen retrieval was conducted by boiling slides in 10 mM sodium citrate buffer (pH 6.0), followed by permeabilization with 1x Tris-Buffered Saline with Tween 20 (TBST). Sections were blocked for 4 hours with blocking buffer (5% bovine serum albumin (BSA) in 1xTBST), then incubated overnight at 4°C with primary antibodies diluted in blocking buffer: rabbit anti-TMEM119 (1:500; Abcam) or rabbit anti-PSD95 (1:1000; GeneTex). On day 2, sections were incubated with Alexa Fluor 555 secondary antibody (1:200; ThermoFisher) diluted in blocking buffer, counterstained with 4′,6-diamidino-2-phenylindole (DAPI;1 μg/mL), and mounted with 10% glycerol in phosphate-buffered saline (PBS). Epifluorescence imaging was performed on a Zeiss Axio Observer Z.1 inverted microscope using a 40× objective and halogen illumination, with Zeiss filter set 49 for DAPI and set 20 for Alexa Fluor 555. Images were processed in ImageJ.

### Functional outcome assessment

A battery of functional assays was conducted 2-4 weeks after the final CHI/sham-injury. Because functional impairments are a common outcome of repetitive mild traumatic brain injury (rmTBI), including depression^4,5^, anxiety^6,46^, and short term memory loss^14,46^, we chose functional assessments that assess these metrics, including tail suspension, open field, and novel object recognition tests. For each testing paradigm, mice were placed in the testing room at least 1 hour prior to testing to acclimate to the environment.

#### Tail suspension test

A tail suspension test (TST) was used to assess depressive-like behavior^47^. Mice were suspended for 6 min at a height of 0.5 m using medical-grade tape placed ∼1 cm from the tip of the tail^48,49^. Immobility time was defined as the summed duration of periods of time for which the animal exhibited passive hanging, absence of body movement, absence of tail-climbing attempts, pendulum motion attributable to prior movement, or movement restricted to either hind or forelimbs. Two blinded researchers scored each video for immobility time, and discrepancies in their scores were adjudicated by a third reviewer to generate a final, multi-researcher vetted total immobility time. This analysis was performed using in-house written software that is publicly available^50^. Greater immobility time was interpreted as depression-like behavior. The apparatus was cleaned with 70% ethanol between mice.

#### Open field test

An open field test was used to assess anxiety-like behavior^51,52^. For the test, mice freely explored a 40 × 40 cm arena for 10 min while an overhead camera recorded their movement. EthoVision software (Noldus) quantified the total time spent in the 20 × 20 cm center zone as well as total mobile time within the 10 min trial. Greater percentage time spent in open field center zone was interpreted as risk-taking or impulsive behavior^51^; greater percentage of mobile time was interpreted as hyperactivity^52^. The arena was cleaned with 70% ethanol between mice.

#### Novel object recognition

The novel object recognition (NOR) test was used to assess memory^53^. The open field test served as habituation to the arena used for NOR. Twenty-four hours after this habituation, an object was placed in the arena and mice were allowed to explore for 10 minutes (Trial 1). Ten minutes after Trial 1 ended, mice were returned to the arena, which now contained the “familiar” object from Trial 1 and a novel object; mice were allowed 10 minutes to explore the arena (Trial 2). For Trial 2, an overhead camera recorded movement, and software (EthoVision, Noldus TX) was used to quantify the investigation time for each object, defined as the total time the mouse spent with their head directed toward the object (defined as a 3 cm radius circle centered at the center of the object) while their head was inside of a 12 x 12 cm zone containing the object. Trials where the mouse did not investigate either object for a summed time of at least 20 seconds were excluded from analysis. A discrimination index was defined as (novel object investigation time − familiar object investigation time) / (novel object investigation time + familiar object investigation time). Higher positive discrimination index value was interpreted as greater preference for the novel object and better short-term memory. The arena and objects were thoroughly cleaned with 70% ethanol between each trial.

### Cerebral blood flow

Non-invasive optical measurements of an index of cerebral blood flow (CBFi) were made with diffuse correlation spectroscopy (DCS) at the following timepoints: baseline (within 24 hours before first injury/sham injury) and 4 hours following the final CHI/sham injury. For these measurements, anesthesia was induced with 3-5% isoflurane (1L/min, 100% oxygen) for ∼2 min and then maintained at 1.5% isoflurane (0.5L/min, 100% oxygen). After allowing the mouse to stabilize for at least 4 minutes, data was acquired (20 Hz) by manually holding a sensor on the dorsal portion of the head for ∼5 seconds (**Fig. S1**). Measurements were repeated 4 times, repositioning slightly between repetitions to account for local inhomogeneities in the interrogated tissue. While the scalp was intact for these measurements, hair was removed with depilatory cream to enhance the signal-to-noise ratio of the detected signal.

DCS data were acquired with a custom built device consisting of an 852 nm long coherence laser (iBeam Smart, TOPTICA Photonics, Farmington, New York, USA), a four-channel single photon counting module (SPCMQA4C-IO, Perkin-Elmer, Quebec, Canada), and a counter/timer data acquisition board (PCIe-6612, National Instruments, Austin, Texas, USA). The DCS sensor contained a source optode (FT-400-EMT, Thorlabs Inc., Newton, New Jersey, USA) and 2 single mode detector optodes (780HP, Thorlabs) spaced 4.5 mm from the source over the right and left hemispheres (**Fig. S1**). DCS-measured intensity autocorrelation functions, g2(t,r,*τ*), at each detector location, r, time, t, and delay time, *τ*, were first down sampled to 4 Hz by averaging to increase signal-to-noise and then fit to the semi-infinite solution to the correlation diffusion equation for CBFi(r,t) with units of cm^2^/s. Fits were constrained to g2>1.05 and assumed fixed absorption and reduction scattering coefficients at 852 nm of 0.25 and 9.35 cm^-1^, respectively, and an index of refraction of 1.4^54^. To ensure high quality data, several quality control metrics were employed: 1) for each repetition and each detector, outlier data frames that fell outside ±1.5 times the standard deviation of the mean CBFi across the repetition were removed, 2) repetitions with mean CBFi that fell outside ±1.5 times the standard deviation of the mean CBFi across all repetitions for a given detector were removed, 3) measurements with a coefficient of variation across all repetitions for a given detector (defined as standard deviation divided by mean) greater than 0.3 were removed. Finally, to quantify a mean regional CBFi for each measurement timepoint, CBFi (r,t) were averaged across all data frames, repetitions, and hemispheres.

### Statistical Analysis

Data were analyzed and figures were generated in RStudio (Boston, MA) using the R programming language. The tidyverse collection of R packages was used for data processing. The R package heatmap3 was used to create heatmaps. The R packages ggplot2, ggpubr, ggbeeswarm, and ggprism were used to generate box plots. The stats::hclust function was used to perform hierarchical clustering using Euclidean distance and the unweighted pair group method with arithmetic mean (UPGMA). For functional, cerebral hemodynamic, and protein analyses, p-values were estimated from Wilcoxon rank sum tests. Bonferroni adjustment was applied to correct for multiple comparisons.

To account for genotype variances observed in all outcome metrics, all data were normalized to the median score of sham-injured animals within each genotype. To account for cohort variance observed in measurements of Tmem119, CD68, and PSD95 from tissues collected 4 weeks post-injury, the limma::RemoveBatchEffect function in R was utilized to conduct computations.

Multivariate cytokine profiles were analyzed using principal component analysis (PCA) to identify principal components (PCs) that represent the greatest variances across groups. Two principal components (PC1 and PC2) were identified, and 5-fold cross-validation was applied to evaluate model robustness. Analyses were performed with the R package ropls. Prior to modeling, cytokine values were z-scored.

## Results

### Neuronal p38ɑ knockout protected against injury-induced functional deficits at 4-weeks post rmTBI

We first investigated whether neuronal p38ɑ drives adverse behavioral outcome after injury. In males, injury increased TST immobility time compared to sham-injury in WT mice (5xCHI + WT vs. sham + WT, p=0.033, **Fig. 1A**). This increase in immobility time was not present in neuronal p38ɑ knockout (CRE) mice (5xCHI + CRE vs. sham + CRE, p=0.6; 5xCHI+ WT vs. 5xCHI+ CRE, p=0.002). These data suggest that neuronal p38α knockout protected against injury-induced depressive-like behavior. Similarly, injury increased the percentage of time spent mobile on the open field test compared to sham-injury in WT animals (5xCHI + WT vs. sham + WT, p=0.036, **Fig. 1B**); this increase was not present in neuronal p38ɑ knockout animals (5xCHI+ CRE vs. sham + CRE, p=0.29; 5xCHI + WT vs. 5xCHI + CRE, p <0.001), suggesting that injury-induced hyperactivity was protected by neuronal p38ɑ knockout. No changes with injury were seen in the time spent in the center zone of the open field test in either the WT or CRE mice (**Fig. 1C**). In addition, injury reduced the discrimination index on NOR in WT mice (5xCHI + WT vs. sham + WT, p=0.034; **Fig. S2**); this decrease was not present in CRE mice (5xCHI+ CRE vs. sham + CRE, p=0.66; 5xCHI + WT vs. 5xCHI + CRE, p=0.16; **Fig. S2**), suggesting that neuronal p38α knockout protected against injury-induced short-term memory loss in males.

**Fig 1.**
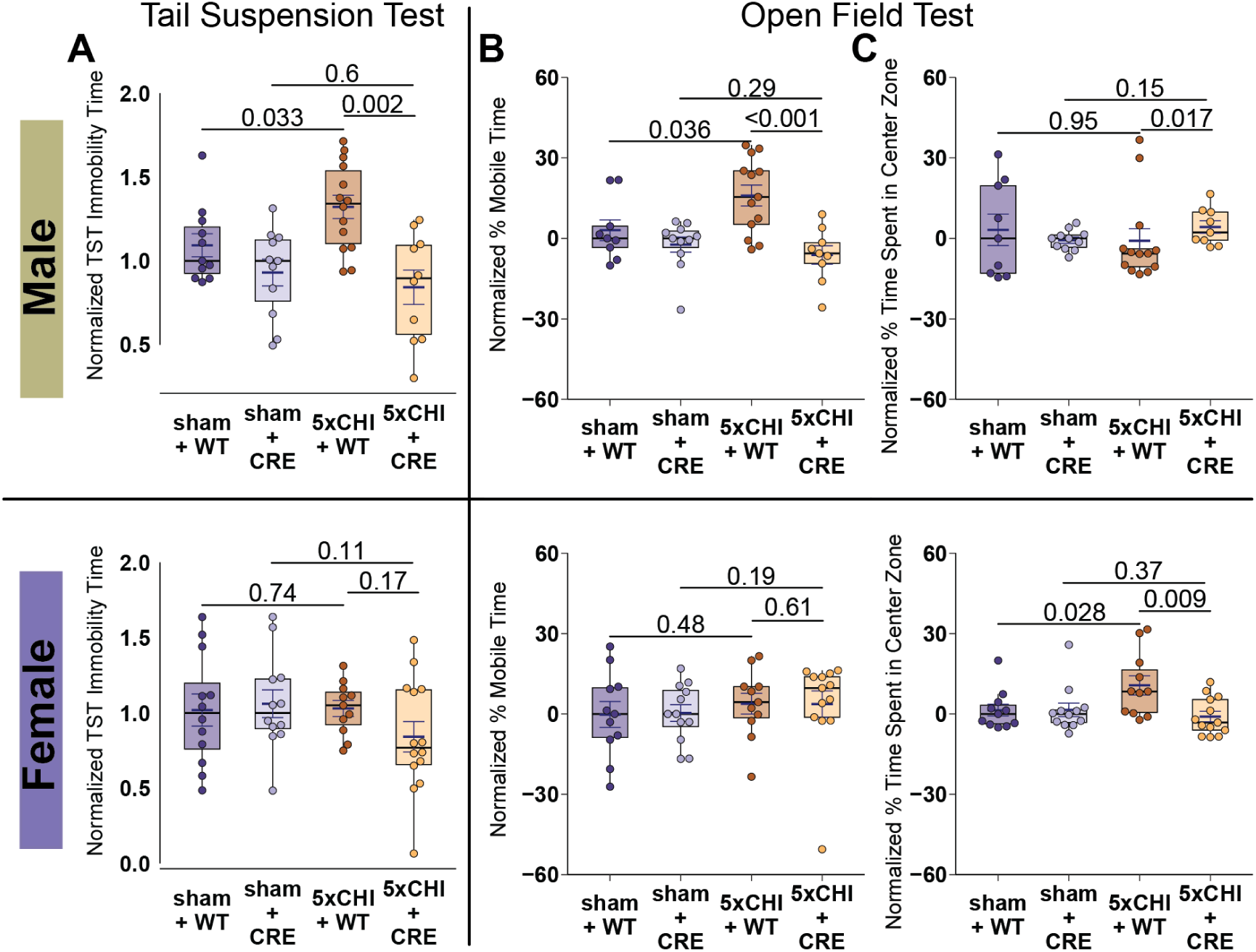
Neuronal p38ɑ knockout animals did not exhibit injury-induced functional deficits at 4 weeks post injury. Box plots of **A.** tail suspension test immobility time (n=11-14/group), **B**. percentage mobile time on open field testing (n=9-13/group), and **C**. percentage of time spent in the center zone on open field testing (n=9-13/group). Males are reported in the top row and females in the bottom row. All data were normalized to the median of the corresponding sham-injured group for each genotype. Each dot denotes one animal. mean±SEM are also displayed. p-values are estimated from Wilcoxon rank sum tests with Bonferroni adjustment for multiple comparisons. 5xCHI = 5 once-daily closed-head injuries; WT = p38ɑ^fl/fl^; wild type control mice; CRE = p38ɑ^fl/fl^; Cre mice.

In females, TST immobility times were not significantly different after injury in either WT or CRE mice, although there was a trend towards decreasing immobility scores after injury in the CRE animals (5xCHI + CRE vs. sham + CRE, p = 0.11, **Fig. 1A**). Similarly, no changes were observed in the percentage of time spent mobile on open field testing with injury in either the WT or CRE mice (**Fig. 1B**). In contrast, injury increased the time spent in open field center zone in WT mice (5xCHI + WT vs. sham + WT, p=0.028, **Fig. 1C**); this increase was not present in neuronal p38ɑ knockout mice (5xCHI + CRE vs. sham + CRE, p=0.37; 5xCHI+ WT vs. 5xCHI+ CRE, p=0.009, **Fig. 1C**), suggesting that neuronal p38α knockout protected against injury-induced risk-taking or impulsive behavior in females. No changes were observed in discrimination index on NOR with injury in either the WT or CRE mice (**Fig. S2**).

Overall, these data suggest that neuronal p38α knockout protects against injury-induced depressive-like behavior, hyperactivity, and short-term memory deficit in males. In contrast, neuronal p38α knockout protects against risk-taking or impulsive behavior in males and females. Considering the sex-based differences in these outcomes, we report the remaining results for males and females separately. Male results are reported in the main text and females in the supplementary text, with a summary section at the end of the Results highlighting key findings.

### Neuronal p38ɑ knockout protected against injury-induced synaptic loss in males 4 weeks post injury

We next examined whether neuronal p38α knockout mitigated rmTBI-induced changes in postsynaptic density protein 95 (PSD95), a key regulator of synaptic plasticity and an established marker of synaptic integrity^55,56^. At 4 weeks post-injury, injured WT animals showed a significant decrease in cortical PSD95 compared with sham-injured controls (5xCHI + WT vs. sham + WT, p=0.007; **Fig. 2A**). This decrease with injury was absent in neuronal p38α knockout animals (5xCHI + CRE vs. sham + CRE, p=0.55; 5xCHI+ WT vs. 5xCHI+ CRE, p=0.019). Immunohistochemistry (IHC) for PSD95 was consistent with these patterns (**Fig. 2B, Fig. S3**). Collectively, these findings indicate that neuronal p38α knockout protects against injury-induced postsynaptic loss at 4 weeks post-injury in males.

**Fig 2.**
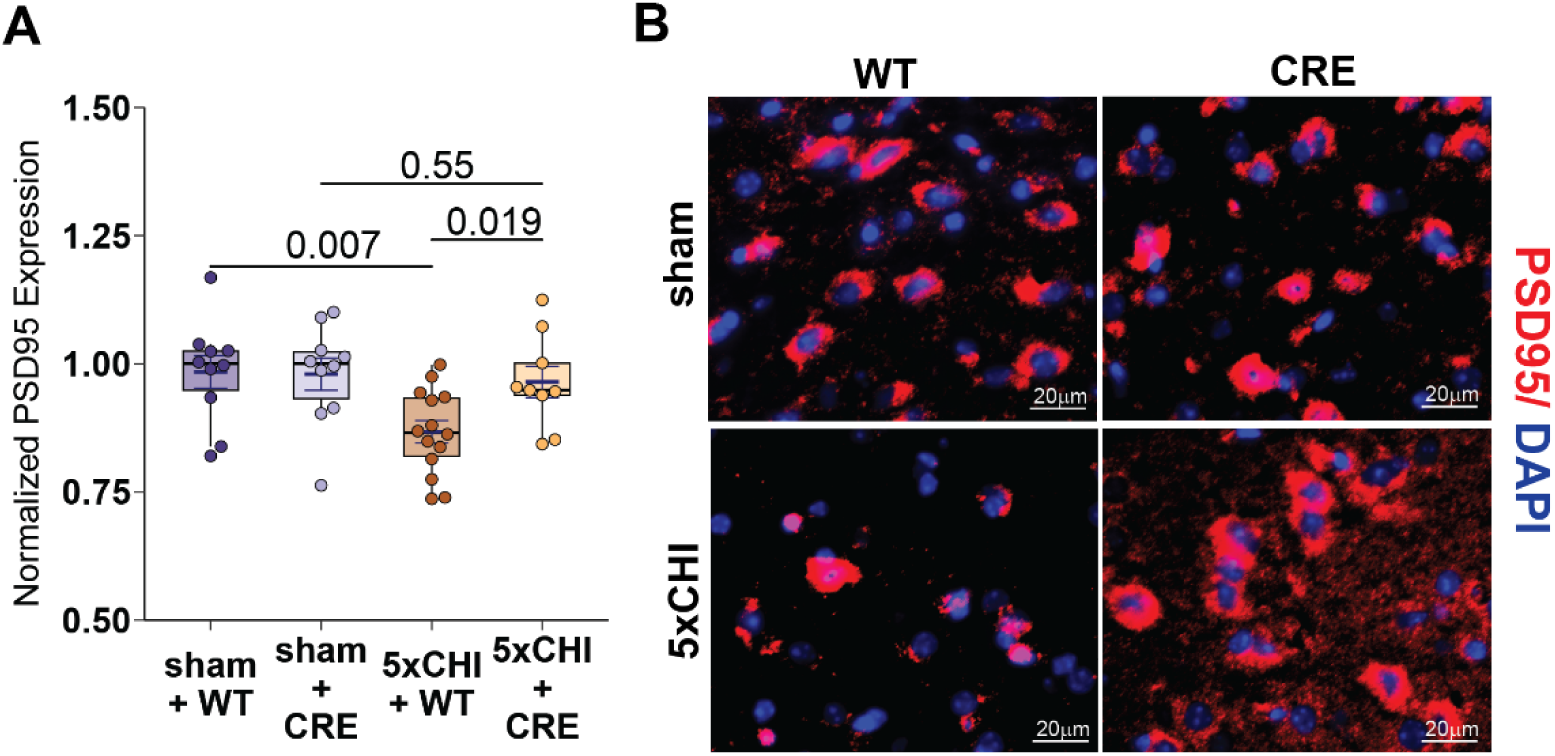
Neuronal p38ɑ knockout animals did not exhibit injury-induced synaptic loss at 4 weeks post injury. **A**. Box plot of PSD95 ELISA results (n=9-14 /group). All values reported were normalized by the means of the corresponding sham-injured group for each genotype. Each dot denotes an individual animal. **B**. Representative immunohistochemistry images in frontal cortex showing PSD95 stain (red) and DAPI (blue) (scale bar: 20µm, representative sections from n=3-4 mice/group). mean±SEM are also displayed. p-values are estimated from Wilcoxon rank sum tests with Bonferroni adjustment for multiple comparisons. PSD95 = post synaptic marker 95; 5xCHI = 5 once-daily closed-head injuries; WT = p38ɑ^fl/fl^; wildtype control mice; CRE = p38ɑ^fl/fl^; Cre mice, DAPI = 4′,6-diamidino-2-phenylindole.

### Neuronal p38ɑ knockout protected against injury-induced microglial changes in males post rmTBI

Given that microglia play a substantial role in the inflammatory cascade post rmTBI^10,12,16,18,26^, we next asked whether neuronal p38ɑ knockout protects against injury-induced microglial changes following rmTBI. At 4 hours post-injury, ELISA of hippocampal samples showed a significant decrease in Tmem119 expression in injured versus sham-injured WT mice (5xCHI + WT vs. sham + WT, p=0.006, **Fig. 3A**), consistent with a decrease in homeostatic microglia. This decrease was not present in neuronal p38ɑ knockout mice (5xCHI + CRE vs. sham + CRE, p=0.85; 5xCHI + CRE vs. 5xCHI + WT, p=0.005). Concordantly, CD68 expression was significantly increased in 5xCHI compared to sham-injured WT mice (5xCHI + WT vs. sham + WT, p=0.027, **Fig. 3B**); this increase was not seen in neuronal p38ɑ knockout mice (5xCHI + CRE vs. sham + CRE p=0.76; 5xCHI + CRE vs. 5xCHI + WT, p=0.007), suggesting a protective effect on acute microglial phagocytic activity. Consistent with ELISA findings, hippocampal IHC revealed amoeboidal microglia with thicker processes in 5xCHI + WT mice, consistent with reactive microglial morphology, whereas all other groups (sham + WT, sham + CRE, 5xCHI + CRE) mice exhibited a ramified microglial morphology with thinner, longer processes, consistent with homeostatic microglial morphology^57^(**Fig. 3C, Fig. S4**). Cortical IHC was qualitatively similar to hippocampal IHC (**Fig. 3D, Fig. S5**).

**Fig 3.**
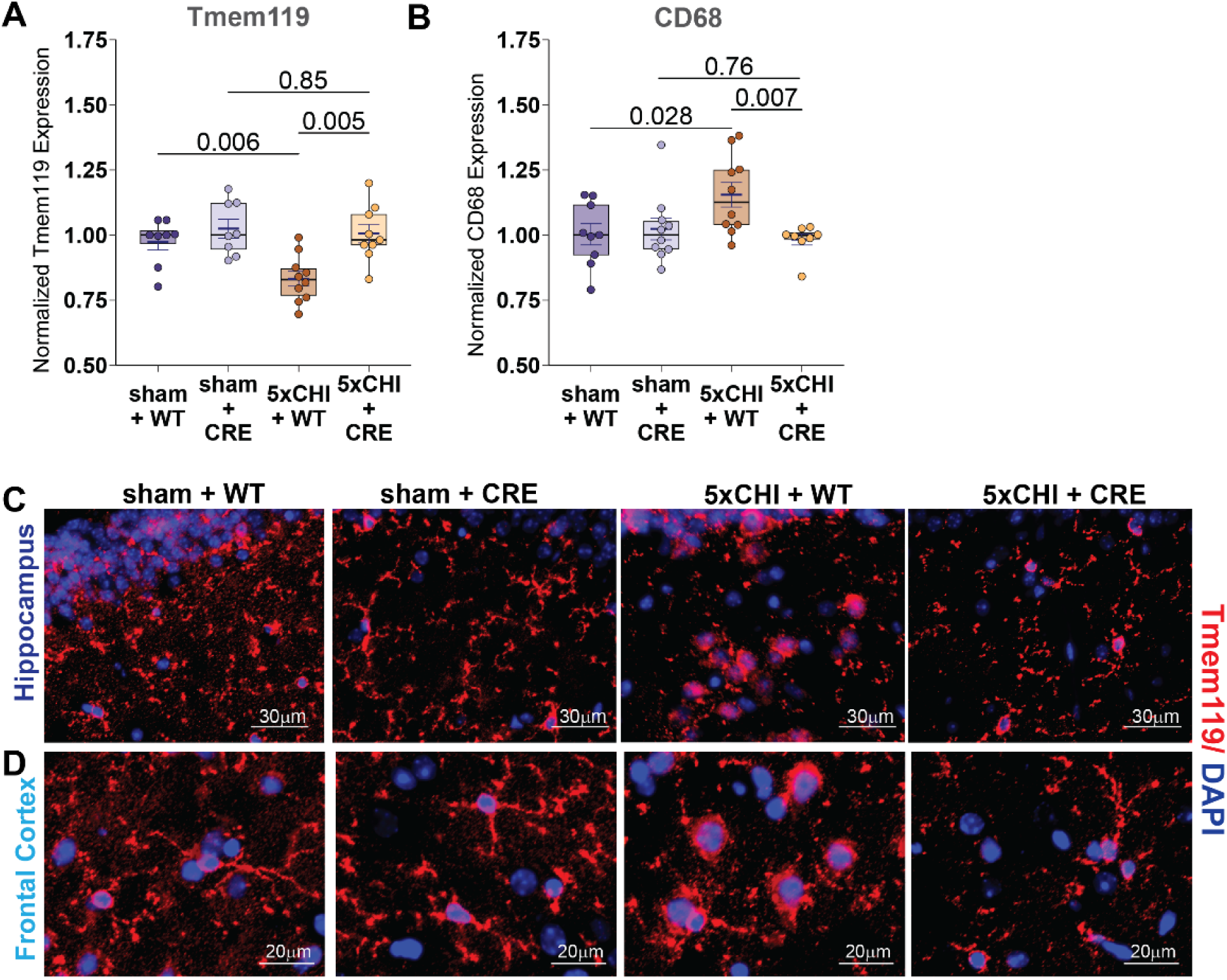
Neuronal p38ɑ knockout animals did not exhibit injury-induced microglial changes post rmTBI in males. Box plots of **A.** Tmem119 ELISA results (n=8-10/group) and **B**. CD68 ELISA results (n=8-10/group). All values reported were normalized by the median of the corresponding sham-injured group for each genotype. Each dot denotes one animal. mean±SEM are also displayed. p-values are estimated from Wilcoxon rank sum tests with Bonferroni adjustment for multiple comparisons. Representative immunohistochemistry images in **C**. hippocampus and **D**. cortex showing Tmem119 stain (red) and DAPI (blue) (scale bar: 30µm, representative sections from n=3-4 mice/group). 5xCHI = 5 once-daily closed-head injuries; WT = p38ɑ^fl/fl^; wildtype control mice; CRE = p38ɑ^fl/fl^; Cre mice. ELISA = enzyme-linked immunosorbent assay, CD68 = Cluster of Differentiation 68, Tmem119 = transmembrane protein 119, DAPI = 4′,6-diamidino-2-phenylindole.

At 4 weeks post-injury, ELISA results demonstrated limited microglial changes with injury in the hippocampus or the frontal cortex (**Fig. S5**). While no changes were seen with TMEM119, cortical CD68 expression was reduced in injured compared to sham-injured CRE mice (5xCHI + CRE vs. sham + CRE, p=0.079, 5xCHI + CRE vs. 5xCHI + WT, p<0.001; **Fig. S5**). Because injury did not significantly change CD68 in WT mice, this genotype difference may be driven by neuronal p38α knockout in the injured condition. Visual inspection of cortical IHC showed similar patterns to those observed at 4 hours post-injury with thicker, amoeboid microglia in 5xCHI + WT and thin, ramified microglia in sham + WT and 5xCHI + CRE mice (**Fig. S6**).

Together, these results demonstrated p38ɑ knockout was protective against microglial changes at 4 hours post rmTBI, with a limited effects persisting through 4 weeks post-injury.

### Neuronal p38ɑ knockout protected against injury-induced inflammatory response in males 4 hours post rmTBI

To assess changes in the inflammatory response post injury, we profiled 18 cytokines via Luminex multiplex assay of hippocampal samples 4 hours post-injury. In males, injury led to an overall upregulation of cytokine expression in WT mice, which was not present in neuronal p38ɑ knockout mice (**Fig. 4A**). Principal component analysis (PCA) identified a first principal component that separated groups and explained 55% of the variance (PC1), while the secondary component (PC2) explained an additional 12% of the variance (**Fig. 4B, Fig. S7A**). PC1 scores indicated that 1) most 5xCHI + WT samples clustered apart from 5xCHI + CRE samples and sham + WT samples, and 2) most 5xCHI + CRE clustered close to sham controls, indicating attenuation of injury effects in injured CRE animals (**Fig. 4B**). Scores on the PC1 were significantly higher in injured versus sham WT animals (5xCHI + WT vs. sham + WT, p=0.002); similar increases in PC1 scores were not present in CRE animals (5xCHI + CRE vs. sham + CRE p = 0.72; 5xCHI + WT vs. 5xCHI + CRE, p = 0.008; **Fig. 4C**). Of the cytokines that contributed to PC1 (**Fig. 4D**), RANTES, TNFɑ, Eotaxin, IL-13, IL-3, M-CSF, IL-2, IP-10, IL-1β, and MIP-1ɑ were significantly elevated with injury in WT mice (all 5xCHI + WT vs. sham + WT p < 0.05) but not in CRE mice (all 5xCHI + CRE vs. sham + CRE p>0.05). This elevation was also reduced in injured CRE mice compared to injured WT mice (5xCHI + WT vs. 5xCHI + CRE, p < 0.05; **Fig. 4E, Fig. S7B, Table S1**). Together, these data demonstrated protective effect of neuronal p38ɑ knockout for injury-induced acute inflammatory response 4 hours post rmTBI.

**Fig 4.**
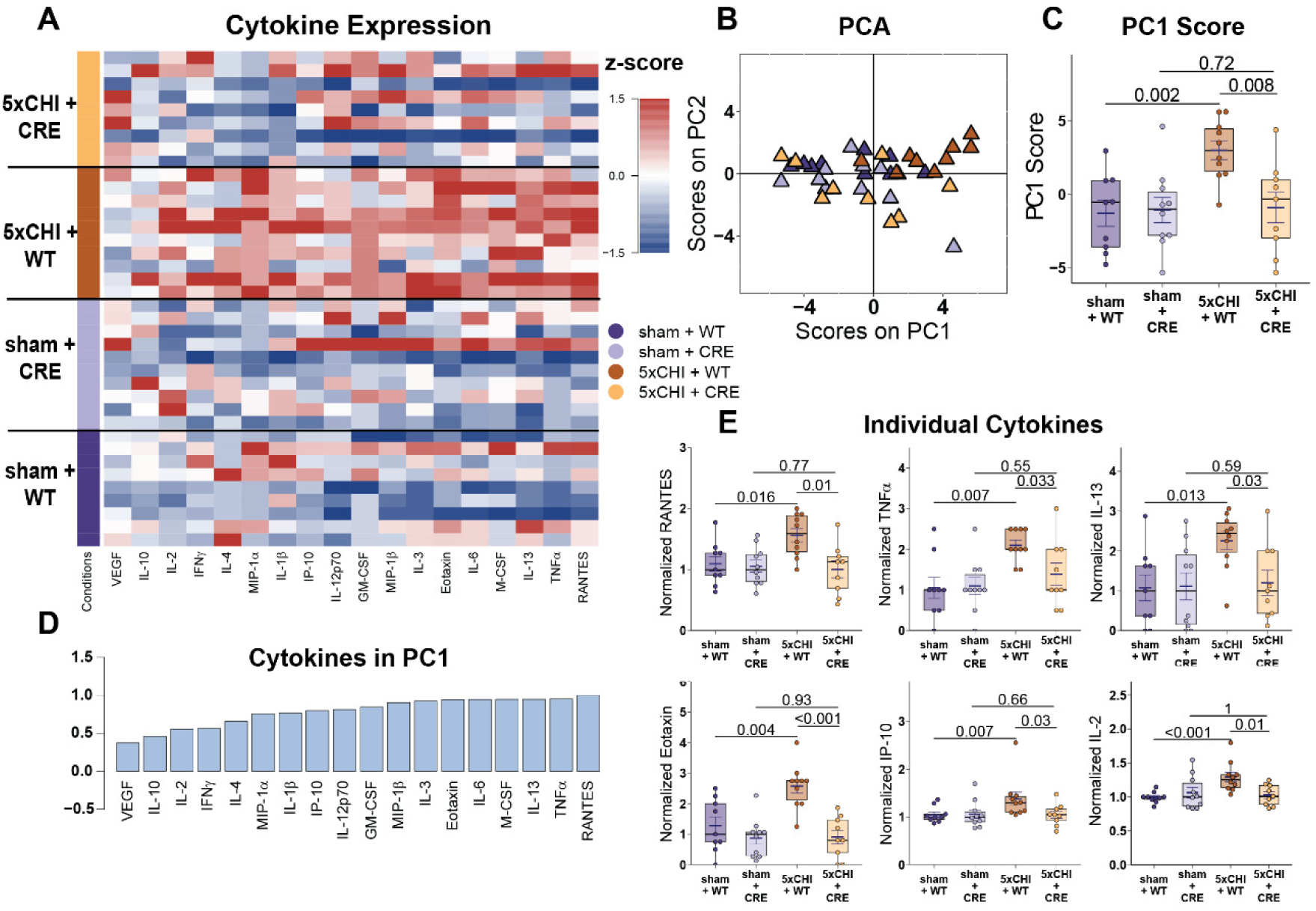
Neuronal p38ɑ knockout animals did not exhibit injury-induced elevation in cytokine expression at 4 hours post rmTBI. **A**. Luminex analysis of 18 hippocampal cytokines at 4 hours after 5xCHI or sham-injury (n=9-10). Each column represents one cytokine analyte, while each row denotes an individual animal (columns are z-scored) **B**. Scores on principal component 1 (PC1) seperated the 5xCHI+WT group to the right. Each triangle denotes one animal. **C**. Boxplot of the scores on PC1 by group (Wilcoxon rank sum tests). **D**. PC1 consisted of a profile of cytokines that were positively correlated with PC1 and thus the 5xCHI+WT group in **C**. **E**. Box plots for individual cytokines that were among the highest weights on PC1. In these plots, all values were normalized by the median of the corresponding sham-injured group for each genotype. Each dot denotes one animal. mean±SEM are also displayed. p-values are estimated from Wilcoxon rank sum tests with Bonferroni adjustment for multiple comparisons. a.u.= arbitrary units. 5xCHI = 5 once-daily closed-head injuries. WT = p38ɑ^fl/fl^; wildtype control mice. CRE = p38ɑ^fl/fl^; Cre mice.

### Neuronal p38ɑ knockout protected against injury-induced reduction of CBF 4 hours post rmTBI

This model has also been shown to reduce cerebral blood flow (CBF^58,59^. Accordingly, we concluded by assessing whether neuronal p38α knockout altered the CBF response to rmTBI. At 4 hours post injury, injured WT animals showed a significant decrease in CBFi compared with sham-injured controls (5xCHI + WT vs. 5xCHI + CRE, p=0.004; **Fig. 5A**); this decrease was not present in injured CRE compared to sham-injured CRE animals (5xCHI + CRE vs. sham + CRE, p=0.64; 5xCHI+ WT vs. 5xCHI+ CRE, p=0.005), indicating that neuronal p38α knockout protects against injury-induced CBF reductions at this acute timepoint.

**Fig. 5.**
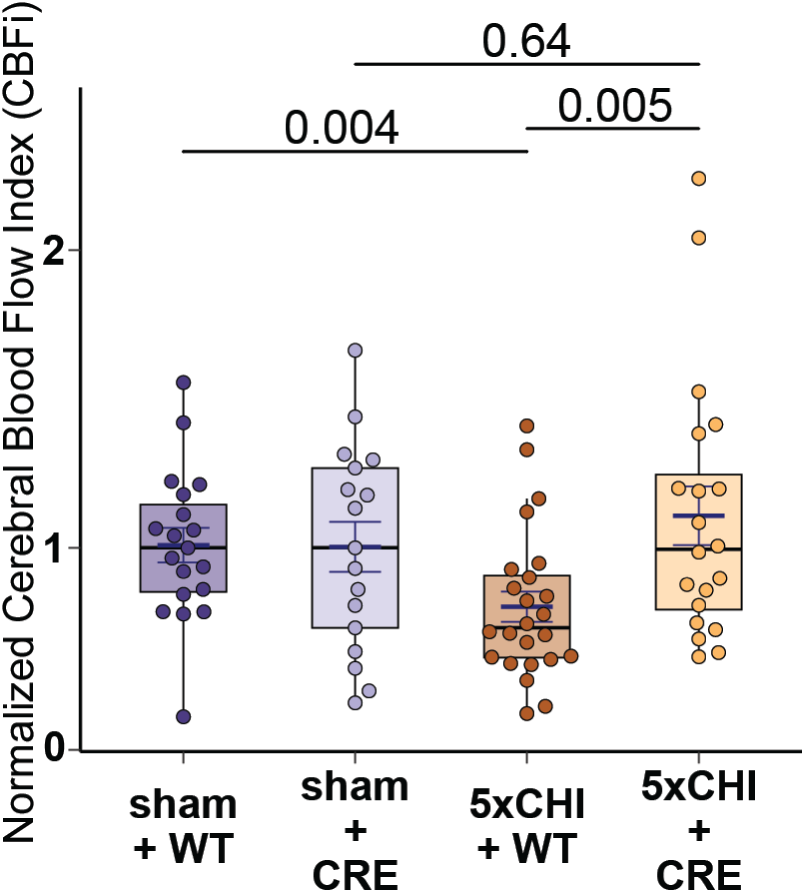
Neuronal p38ɑ knockout animals did not exhibit injury-induced decreases in cerebral blood flow (CBF). **A**. Box plot of the normalized cerebral blood flow index (CBFi) at 4 hours after 5xCHI or sham-injury dichotomized by group (n=17-24/group). All values were normalized by the median of the corresponding sham-injured group for each genotype. Each dot denotes one animal. mean±SEM are also displayed. p-values are estimated from Wilcoxon rank sum tests with Bonferroni adjustment for multiple comparisons. 5xCHI = 5 once-daily closed-head injuries; WT = p38ɑ^fl/fl^; wildtype control mice; CRE = p38ɑ^fl/fl^; Cre mice.

### Neuronal p38ɑ knockout had limited protective effects against the acute and long-term consequences of rmTBI in females

Overall, females exhibited a less pronounced response to injury than males. Among the measures that did demonstrate an injury effect in WT females, neuronal p38α knockout conferred a comparatively weaker protective effect than that observed in males (summarized in **Fig. 6**).

**Fig. 6.**
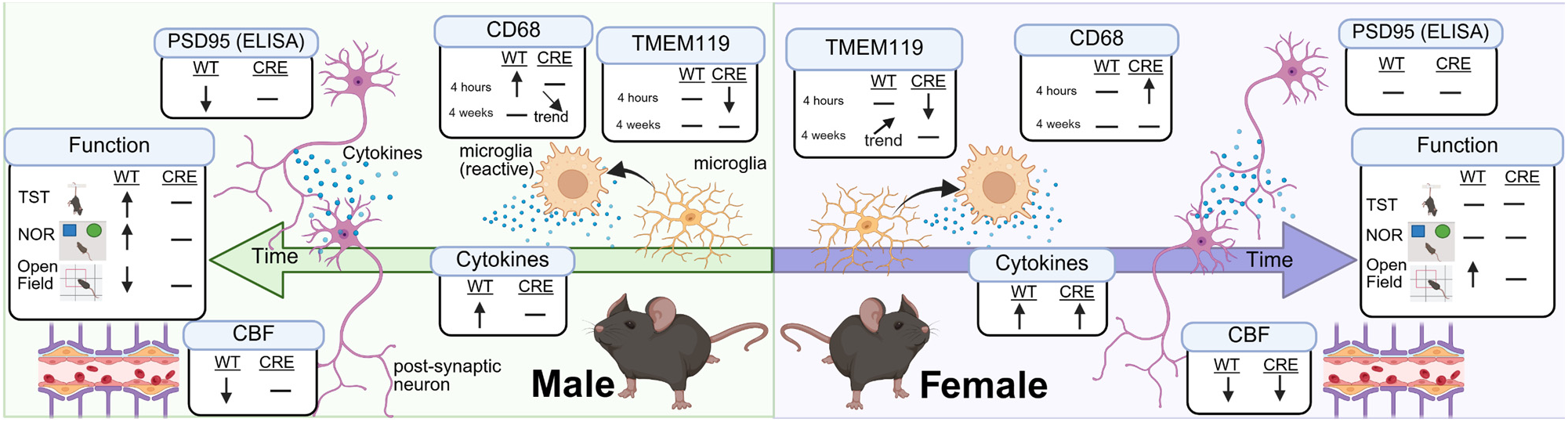
Summary of injury responses in WT and CRE mice across females and males. Summary of responses for male and female mice across major outcome metrics. WT indiates injury response in WT mice; CRE indicates injury response in CRE mice. An upward arrow indicates a significant increase (p<0.05), a downward arrow indicates a significant decrease (p<0.05), and horizontal dash indicates little or no change. Angled arrows denote trends (p<0.2) that did not reach statistical significance (p>0.05). Created in BioRender. Rivera Moctezuma, F. (2026) https://BioRender.com/zm9la7e.

For PSD95, females exhibited a non-significant trend toward injury-induced synapse loss in WT mice (5xCHI + WT vs. sham + WT, p = 0.2; **Fig. S8A**). This decrease was not seen in injured CRE (neuronal p38α KO) animals compared to sham-injured CRE animals (5xCHI + CRE vs. sham + CRE, p=0.97). Qualitative immunohistochemistry (IHC) of PSD95 confirmed these patterns (**Fig. S8B**). Together, these data suggested that injury had minimal effects on synaptic loss in both WT and CRE animals.

For glial markers, WT females showed no response to injury at 4 hours post injury. In contrast, CRE females had decreased hippocampal Tmem119 expression after injury and WT females had increased CD68 expression after injury (**Fig. S9**). This difference was not visually apparent in IHC labeling for Tmem119, potentially because the subtle changes detected by ELISA were not resolved at the tissue level with immunolabeling and/or because ELISA quantifies total protein abundance while IHC primarily reflects cell-level localization and morphology (**Fig. S10-13**). At 4 weeks post injury, injury tended to increase cortical Tmem119 expression in injured WT compared to sham WT mice, although this result was not statistically significant (5xCHI + WT vs. sham + WT, p = 0.075; **Fig. S11**). This increase was absent in injured CRE mice (5xCHI + CRE vs. sham + CRE, p=0.41; 5xCHI + WT vs. 5xCHI + CRE, p = 0.036; **Fig. S11**). These findings were consistent with visual inspection of IHC wherein injured CRE mice exhibited thin processes and a ramified microglial pattern similar to that of sham-injured CRE and WT mice, whereas injured WT mice exhibited thicker, denser processes (**Fig. S12**). Taken together, these results indicate that 1) neuronal p38α knockout amplified injury response at 4 hours post injury, and 2) neuronal p38α knockout protected against injury-induced microglial changes in females at 4 weeks post injury.

For acute cytokine response at 4 hours after the final injury, female hippocampal cytokine profiles showed pronounced injury effects in both WT and CRE mice, with limited effects of neuronal p38α knockout (**Fig. S13**). Of the cytokines measured, VEGF, MIP-1β, M-CSF, Eotaxin, IL-12p70, MIP-1ɑ, IP-10, and IL-3 were significantly upregulated by injury in WT mice (5xCHI + WT vs. sham + WT, p < 0.05; **Fig. S13F; Table S2**). Among these, VEGF and MIP-1ɑ did not change with injury in CRE mice (5xCHI + CRE vs. sham + CRE, p>0.05). Although injury significantly increased M-CSF, Eotaxin, and IL-12p70 in CRE mice (5xCHI + CRE vs. sham + CRE, p<0.05), these increases with injury were less pronounced in neuronal p38ɑ knockout CRE mice compared to WT mice. Together, these findings indicate that females mounted a robust acute cytokine response to injury in both WT and CRE mice; however, this increase was attenuated in a subset of cytokines in CRE compared to WT mice.

Finally, injury significantly decreased CBF at 4 hours post injury in both female WT and female CRE mice (5xCHI + WT vs. sham + WT, p = 0.014; 5xCHI + CRE vs. sham + CRE, p= 0.033; **Fig. S14**). No differences in the magnitude of these decreases were observed between the WT and CRE injured groups (5xCHI + WT vs. 5xCHI + CRE, p=0.66), suggesting a pronounced effect of injury on cerebral hemodynamics that is not protected by by neuronal p38α knockout in females.

Taken together, these data suggest that neuronal p38ɑ knockout in females partially protects against injury-induced changes, including time spent in open field center zone, longer-term microglial changes, and MIP-1β expression, while having limited effects on overall cytokine expression, PSD95 loss, and reductions in CBF.

## Discussion

Herein we demonstrate that neuronal p38α is a mediator of the functional, immune, and cerebrovascular consequences of repetitive mild traumatic brain injury (rmTBI), establishing a cell type–specific causal mechanism for rmTBI-associated dysfunction. To our knowledge, this is the first work to examine neuronal p38α in a mild, repetitive TBI model.

As expected based on previous reports from both mouse^60–62^ and human data^6,31,63–65^, we observed substantial sex-dependent injury differences in the majority of outcome metrics that we investigated. For example, male WT mice showed injury-induced depressive-like behavior in TST (**Fig. 1A**), short term memory deficit<u>s</u> in NOR (**Fig. S2**), significant reductions in PSD95 at 4 weeks post-rmTBI (**Fig. 2**), and acute changes in microglial markers (**Fig. 3**), whereas WT females showed no change in these parameters (**Fig. 1A; Fig. S10**). Possible explanations for these sex-based differences include sex-dependent injury kinetics, potentially arising from differences in baseline immune profile and/or post-injury immune timing^18^. Notably, while we found more prominent injury-induced changes in TST, PDS95 and microglial markers in males herein, prior work using this same injury model and sampling timepoints found these injury effects were more prominent in females^31^. This discrepancy may be caused by several factors. First, although we included a two-week washout period, tamoxifen can be immunomodulatory^66,67^ and has been shown to alter baseline behavioral outcomes in other CreERT2 lines (e.g., CaMKIIα-CreERT2 and Aldh1l1- CreERT2)^68^, potentially altering baseline behavior and immune profiles, and thereby influencing the post-injury response. Alternatively, despite an otherwise WT background, sex-dependent variation in baseline Cre activity has been reported^69^ and could potentially shift baseline immunity and behavior, thereby modulating the injury response.

We also observed sex-specific differences in the effects of neuronal p38α knockout on the injury response. In males, injury-induced cytokine upregulation observed in WT mice was absent in injured CRE mice, whereas this attenuation was less evident in females (**Fig. 4; Fig. S13**). Similarly, WT males had significant decreases in CBF after injury while CRE males did not exhibit changes in CBF after injury; similar absence of CBF changes after injury were not observed in female CRE mice (**Fig. 5; Fig. S14**). These sex-specific modulations suggest that the contribution of neuronal p38α to downstream injury consequences may differ between males and females, supporting the notion of sex-distinct causal mechanisms following repetitive mild TBI.

This injury model induces clinically-relevant functional deficits, including depressive-like behavior, hyperactivity, and short-term memory impairment (**Fig. 1; Fig. S2**), all of which are commonly reported after mTBI/rmTBI^46^. In males, TST immobility times, open field mobility times, and NOR discrimination index were changed with injury in WT but not in CRE mice, suggesting protection against depressive-like behavior, anxiety-like behavior, and short-term memory deficits. Together with broader evidence linking p38 MAPK signaling to post-injury dysfunction across TBI models^27,28,31,70^, our results suggests neuronal p38α as a key mediator of post-injury functional changes. Such implication is further supported by prior work demonstrating that neuronal p38α signaling modulates behavior in uninjured animals^71,72^ and in mouse models of Alzheimer’s disease^37^. For example, selective deletion of p38α in serotonergic neurons reduces depressive-like behavior in stress-related paradigms (social defeat stress–induced social avoidance and reinstatement of cocaine preference)^71^, and pan-neuronal p38α deletion modulates anxiety-related behavior^72^, indicating that neuronal p38α signaling can modulate functional outcomes across baseline and injury contexts.

In parallel with functional protection, injury-induced PSD95 loss was absent in male injured CRE mice (**Fig. 2; Fig. S3**), supporting a role for neuronal p38α in injury-induced synaptic loss and strengthening the link between neuronal p38α activity and downstream functional outcomes. Given that reduction of PSD95 has been associated with compromised synaptic function and cognitive decline^55,73–75^, preservation of PSD95 provides a plausible route for functional protection. Mechanistically, neuronal p38α may promote synaptic loss after rmTBI either through direct synaptotoxic effects and/or by initiating inflammatory signaling, consistent with prior links between p38 MAPK activity and synaptic integrity^37,41,76,77^. This function and protein data elucidate a role for neuronal p38α in driving post-injury synaptic loss for the first time, suggesting that that neuronal p38α signaling may drive synapses loss and thereby functional deficits following rmTBI in males. Importantly, this cell-type-specific mechanism builds on our prior pharmacological inhibition study, which demonstrated that p38 MAPK inhibition with the small molecule inhibitor SB239063 ameliorated injury-induced deficits, and specifically implicated the p38α isoform signaling in neurons. Given that depressive symptoms and measurable cognitive impairment are commonly reported after mTBI/rmTBI^46^, these data strengthen the translational rationale for targeting the p38 pathway, specifically neuronal p38α, to protect against synaptic loss and long-term functional consequences following repetitive head injury.

Another key finding is that injury-induced cytokine upregulation at 4 hours after 5xCHI in WT male mice was largely absent in neuronal p38α knockout mice, suggesting neuronal p38α may drive the acute inflammatory response (**Fig. 4; Fig. S7**). This result aligns with our prior observations that several injury-elevated cytokines co-localize with the neuronal marker NeuN^18,59^ and with our previous work showing that inhibition of MAPK ameliorated cytokine upregulation after 5xCHI^31^. Notably, multiple cytokines that were elevated in WT but not CRE male mice, including TNFα^59,78,79^, IL-1β^80,81^, IP-10^82^, RANTES^83,84^, Eotaxin^85^, IL-2^86^, and MIP-1α^87,88^, have been implicated in acute inflammatory cascades and downstream consequences after mTBI/rmTBI, underscoring the phenotypic relevance of their attenuation with neuronal p38α knockout. Although cytokines were measured in whole-tissue lysates rather than neuron-specific fractions, these results are still indicative that restricting p38α signaling in neurons is sufficient to modulate the early inflammatory response, and they motivate future cell-type-resolved signaling proteomics. In parallel, injury-induced increases in CD68 and reductions in Tmem119 observed in WT male mice were absent in CRE males, indicating protection by neuronal p38α knockout against early microglial reactivity. Given the temporal alignment between cytokine upregulation and microglial changes, it is plausible that the reduced microglial reactivity is mediated, at least in part, by diminished cytokine production that resulted from diminished signaling downstream of neuronal p38α phospho-activation. These data, however, do not exclude the potential role of concurrent alterations in microglial p38 MAPK signaling after rmTBI, as previously reported^11^.

We also found that injury-induced decreases in cerebral blood flow (CBF) observed in male WT mice at 4 hours post-5xCHI were absent in male neuronal p38α knockout mice (**Fig. 5**), supporting a protective role for neuronal p38α deletion against early hemodynamic dysfunction. This result is consistent with prior work showing that reduced CBF after rmTBI correlates with, increased monocyte reactivity marker, Iba1, and increased p38 MAPK expression^59^. Because oxidative stress-induced cerebrovascular dysfunction can modulate CBF^89–91^, and neuronal p38α has been implicated in oxidative stress^92,93^, these data suggest that neuronal p38α may contribute to post-injury hemodynamic dysregulation. More broadly, this data aligns with literature implicating p38 MAPK signaling in cerebrovascular regulation in general^94–96^ and in fluid percussion brain injury and subarachnoid hemorrhage models^97,98^. Sankar *et al.* (2019) reported correlations between CBF and cytokines such as IP-10, RANTES, and IL-13 in injured WT animals^59^. Given that neuronal p38α knockout protected against injury-induced upregulation of these cytokines in our study, it is also plausible that preserved CBF was partially mediated by reduced inflammatory signaling. Overall, these findings support a potential role of neuronal p38ɑ in the regulation of cerebral hemodynamics post rmTBI.

This study has several limitations that motivate future work. First, protein markers were quantified from whole-brain lysates, which capture how neuronal p38α knockout modulates overall tissue-level responses but do not resolve cell-type–specific immune profiles. Future studies could apply cell-type specific approaches, such as native-state proteomic labeling, to define neuronal- and glial-specific signaling changes after rmTBI. Second, robust sex differences were observed across nearly all outcome metrics, which may reflect injury biology as well as experimental factors such as the specific time points sampled. Thus, systematically testing the contribution of these parameters will be important in the future. Third, because sham-injured WT and CRE mice often differed at baseline, we normalized outcomes to the median of the sham group within each genotype. This approach helps isolate injury-associated changes, but it also suggests that neuronal p38α deletion (and/or the tamoxifen inducible Cre-lox paradigm) may alter baseline immunity and behavior. Thus, the response to injury reported in this study should be interpreted in the context of these genotype-dependent baseline differences. Fourth, estrous cycle stage was not monitored in female mice. Although emerging evidence suggests estrous-related variability may have modest effects on injury outcomes^99,100^, some assays (e.g., TST, open field) can be hormone sensitive, and incorporating cycle tracking in the future could better delineate hormone-dependent behavioral and molecular responses. Fifth, although we detected CBF changes at 4 hours post-injury, we did not directly assess vascular structure or function because defining vascular mechanisms was not a primary focus of the present study. Given that vascular changes can modulate CBF^89,90,101^ and contribute more broadly to rmTBI pathophysiology^101–103^, future studies could elucidate structural and functional vascular changes (e.g., vessel density, area fraction, vascular reactivity) in neuronal p38α knockout mice to better define the mechanisms linking neuronal p38α to hemodynamic outcomes. Sixth, we assessed only two post-injury timepoints. Given that injury-response patterns in the present study differ from our prior inhibition study^31^, neuronal p38α deletion may alter response kinetics such that key changes occur outside our sampling windows. Future studies should include additional acute and chronic time points (e.g., 24 hours and 3 months post-injury) to better define the temporal profile of these effects. Finally, while we profiled multiple protein markers, we did not perform transcriptomic analysis such as bulk RNA sequencing. Transcriptomic analyses would provide a more comprehensive view of cellular function and pathway-level changes and would help to identify additional protective mechanisms and therapeutic targets modulated by neuronal p38α following rmTBI.

Altogether, this work builds on prior studies implicating p38 MAPK signaling in TBI/mTBI outcomes^11,26–28,31,70^, but uniquely provides evidence for a neuron-specific, causal contribution to post-injury deficits. Given the protection in multiple functional, synaptic, and inflammatory outcome metrics, the data support further investigation into the therapeutic potential of targeting p38α signaling after rmTBI. Importantly, demonstrating that neuronal p38α deletion alone is sufficient to protect against injury-induced dysfunction offers a clear direction for p38-based intervention strategies. Given that p38α is an essential phosphoprotein with diverse homeostatic roles, refining therapies toward greater cell-type and pathway specificity, while preserving efficacy, may enable more precise and potentially safer treatment approaches.

## Acknowledgements

We thank the Wood and Buckley laboratories for critical feedback and technical assistance for this work. We wish to acknowledge Aqua Asberry and David J. Alexander from the histology core facility the Parker H. Petit Institute for Bioengineering and Bioscience at the Georgia Institute of Technology for use of their shared equipment, services, and expertise. We wish to acknowledge Eric Liu for trouble shooting through the immunohistochemistry protocol. We also wish to acknowledge fruitful discussions with Alyssa Pybus, Sara Bitarafan, Yibo Fu, Brendan Tobin, Andrew Feola, and Srikant Rangaraju.

## Ethics approval

All animal protocols were approved by the Emory University Institutional Animal Care and Use Review Office.

## Consent to publish declaration

Not applicable.

## Funding

This work was supported by the National Institutes of Health under Award Nos. 1 R01 NS115994 (LBW/EMB) and by support from the George W. Woodruff School of Mechanical Engineering Faculty Fellowship at Georgia Tech (LBW).

## Contributions

CL conducted data analysis and wrote the manuscript. CL, SET, and FGM prepared figures. SET, AK, PFS, PIS, and CL conducted animal experiments and curated animal data. MNG, CL, ALH, SJS, JCL, RW, SEL, DD, and MC conducted molecular assays and experimentation. LBW and EMB conceived the study and supervised the research. MNG, SET, FGM, LBW and EMB and revised the manuscript. All authors reviewed the manuscript.

## Competing interest

All authors declare that they have no competing interests.

## Notes

### Competing Interest Statement

The authors have declared no competing interest.

## Reference

1. Silverberg, N. D. & Iverson, G. L. Etiology of the post-concussion syndrome: Physiogenesis and psychogenesis revisited. NeuroRehabilitation 29, 317–329 (2011).

2. US Department of Health & Human Services; Centers for Disease Control (CDC); National Center for Injury Prevention and Control. Report to Congress on Mild Traumatic Brain Injury in the United States: Steps to Prevent a Serious Public Health Problem: (371602004-001). 10.1037/e371602004-001 (2003).

3. Langlois, L. D. et al. Repetitive mild traumatic brain injury induces persistent alterations in spontaneous synaptic activity of hippocampal CA1 pyramidal neurons. IBRO Neurosci. Rep. 12, 157–162 (2022).

4. Guskiewicz, K. M. Cumulative effects associated with recurrent concussion in collegiate football players: the NCAA Concussion Study. Jama 290, 2549–55 (2003).

5. Wilk, J. E., Herrell, R. K., Wynn, G. H., Riviere, L. A. & Hoge, C. W. Mild traumatic brain injury (concussion), posttraumatic stress disorder, and depression in U.S. soldiers involved in combat deployments: association with postdeployment symptoms. Psychosom Med Apr;74(3):249–257. **PMID**, 22366583 (2012).

6. Lumba-Brown, A. et al. Multicentre evaluation of anxiety and mood among collegiate student athletes with concussion. BMJ Open Sport Exerc. Med. 9, e001446 (2023).

7. Longhi, L. Temporal window of vulnerability to repetitive experimental concussive brain injury. Neurosurgery 56, 364–374 364-374 (2005).

8. Lennon, M. J. et al. Lifetime Traumatic Brain Injury and Cognitive Domain Deficits in Late Life: The PROTECT-TBI Cohort Study. J. Neurotrauma 40, 1423–1435 (2023).

9. Meehan, W. P., Zhang, J., Mannix, R. & Whalen, M. J. Increasing recovery time between injuries improves cognitive outcome after repetitive mild concussive brain injuries in mice. Neurosurgery Oct;71(4):885–891, (2012).

10. Ramlackhansingh, A. F. et al. Inflammation after trauma: Microglial activation and traumatic brain injury. Ann. Neurol. 70, 374–383 (2011).

11. Morganti, J. M., Goulding, D. S. & Van Eldik, L. J. Deletion of p38α MAPK in microglia blunts trauma-induced inflammatory responses in mice. J. Neuroinflammation 16, 98 (2019).

12. Bray, C. E. et al. Chronic Cortical Inflammation, Cognitive Impairment, and Immune Reactivity Associated with Diffuse Brain Injury Are Ameliorated by Forced Turnover of Microglia. J. Neurosci. Off. J. Soc. Neurosci. 42, 4215–4228 (2022).

13. Ertürk, A. et al. Interfering with the Chronic Immune Response Rescues Chronic Degeneration After Traumatic Brain Injury. J. Neurosci. 36, 9962–9975 (2016).

14. Hu, T. et al. Inhibition of Exosome Release Alleviates Cognitive Impairment After Repetitive Mild Traumatic Brain Injury. Front. Cell. Neurosci. 16, (2022).

15. Lu, S. et al. Decoupling the mutual promotion of inflammation and oxidative stress mitigates cognitive decline and depression-like behavior in rmTBI mice by promoting myelin renewal and neuronal survival. Biomed. Pharmacother. 173, 116419 (2024).

16. Jia, Z.-X. et al. Decreased IL-33 in the brain following repetitive mild traumatic brain injury contributes to cognitive impairment by inhibiting microglial phagocytosis. Mil. Med. Res. 12, 46 (2025).

17. Giza, C. C. & Hovda, D. A. The new neurometabolic cascade of concussion. Neurosurgery 75 **Suppl 4**, S24–33 (2014).

18. Pybus, A. F. et al. Profiling the neuroimmune cascade in 3xTg-AD mice exposed to successive mild traumatic brain injuries. J. Neuroinflammation 21, 156 (2024).

19. Vagnozzi, R. et al. Temporal window of metabolic brain vulnerability to concussions: mitochondrial-related impairment–part I. Neurosurgery Aug;61(2):379–388, 388–389 (2007).

20. Barkhoudarian, G., Hovda, D. A. & Giza, C. C. The Molecular Pathophysiology of Concussive Brain Injury - an Update. Phys Med Rehabil Clin N Am May;27(2):373–93. **PMID**, 27154851 (2016).

21. Wang, Y. et al. Early posttraumatic CSF1R inhibition via PLX3397 leads to time- and sex-dependent effects on inflammation and neuronal maintenance after traumatic brain injury in mice. Brain. Behav. Immun. 106, 49–66 (2022).

22. Henry, R. J. et al. Microglial Depletion with CSF1R Inhibitor During Chronic Phase of Experimental Traumatic Brain Injury Reduces Neurodegeneration and Neurological Deficits. J. Neurosci. Off. J. Soc. Neurosci. 40, 2960–2974 (2020).

23. Wood, L. B. & Singer, A. C. Neurons as Immunomodulators: From Rapid Neural Activity to Prolonged Regulation of Cytokines and Microglia. https://doi.org/10.1146/annurev-bioeng-110122-120158 (2025) doi:10.1146/annurev-bioeng-110122-120158.

24. Pósfai, B., Cserép, C., Orsolits, B. & Dénes, Á. New Insights into Microglia–Neuron Interactions: A Neuron’s Perspective. Neuroscience 405, 103–117 (2019).

25. Miyamoto, A., Wake, H., Moorhouse, A. J. & Nabekura, J. Microglia and synapse interactions: fine tuning neural circuits and candidate molecules. Front. Cell. Neurosci. 7, (2013).

26. Bachstetter, A. D. et al. The p38α MAPK Regulates Microglial Responsiveness to Diffuse Traumatic Brain Injury. J. Neurosci. 33, 6143–6153 (2013).

27. Zhao, X.-Y., Yu, D., Shi, X., Hou, S. & Teng, D. Resveratrol reduces p38 mitogen-activated protein kinase phosphorylation by activating Sirtuin 1 to alleviate cognitive dysfunction after traumatic brain injury in mice. NeuroReport 33, 463 (2022).

28. Zhao, T. et al. Inhibition of TREM-1 alleviates neuroinflammation by modulating microglial polarization via SYK/p38MAPK signaling pathway after traumatic brain injury. Brain Res. 1834, 148907 (2024).

29. Lan, Y.-L. et al. Bazedoxifene protects cerebral autoregulation after traumatic brain injury and attenuates impairments in blood–brain barrier damage: involvement of anti-inflammatory pathways by blocking MAPK signaling. Inflamm. Res. 68, 311–323 (2019).

30. Yan, J. et al. TREM2 activation alleviates neural damage via Akt/CREB/BDNF signalling after traumatic brain injury in mice. J. Neuroinflammation 19, 289 (2022).

31. Li, C. et al. Inhibition of p38 MAPK after repetitive mTBI ameliorates immune signaling and functional deficits. Acta Neuropathol. Commun. https://doi.org/10.1186/s40478-026-02226-w (2026) doi:10.1186/s40478-026-02226-w.

32. Cuadrado, A. & Nebreda, A. R. Mechanisms and functions of p38 MAPK signalling. Biochem. J. 429, 403–417 (2010).

33. Asih, P. R. et al. Functions of p38 MAP Kinases in the Central Nervous System. Front. Mol. Neurosci. 13, (2020).

34. Hou, K., Xiao, Z. & Dai, H.-L. p38 MAPK Endogenous Inhibition Improves Neurological Deficits in Global Cerebral Ischemia/Reperfusion Mice. Neural Plast. 2022, 3300327 (2022).

35. Falcicchia, C., Tozzi, F., Arancio, O., Watterson, D. M. & Origlia, N. Involvement of p38 MAPK in Synaptic Function and Dysfunction. Int. J. Mol. Sci. 21, 5624 (2020).

36. Falcicchia, C., Tozzi, F., Arancio, O., Watterson, D. M. & Origlia, N. Involvement of p38 MAPK in Synaptic Function and Dysfunction. Int. J. Mol. Sci. 21, 5624 (2020).

37. Colié, S. et al. Neuronal p38α mediates synaptic and cognitive dysfunction in an Alzheimer’s mouse model by controlling β-amyloid production. Sci. Rep. 7, 45306 (2017).

38. Li, S. et al. Interleukin-13 and its receptor are synaptic proteins involved in plasticity and neuroprotection. Nat. Commun. 14, 200 (2023).

39. Di Cesare Mannelli, L., et al. Neuronal alarmin IL-1α evokes astrocyte-mediated protective signals: Effectiveness in chemotherapy-induced neuropathic pain. Neurobiol. Dis. 168, 105716 (2022).

40. Canovas, B. & Nebreda, A. R. Diversity and versatility of p38 kinase signalling in health and disease. Nat. Rev. Mol. Cell Biol. 22, 346–366 (2021).

41. Iba, M. et al. Inhibition of p38α MAPK restores neuronal p38γ MAPK and ameliorates synaptic degeneration in a mouse model of DLB/PD. Sci. Transl. Med. 15, eabq6089 (2023).

42. Brothers, R. O., Bitarafan, S., Pybus, A. F., Wood, L. B. & Buckley, E. M. Systems Analysis of the Neuroinflammatory and Hemodynamic Response to Traumatic Brain Injury. J. Vis. Exp. JoVE https://doi.org/10.3791/61504 (2022) doi:10.3791/61504.

43. Mannix, R. et al. Clinical correlates in an experimental model of repetitive mild brain injury. Ann Neurol Jul;74(1):65–75. **PMID**, 23922306 (2013).

44. Korzhevskii, D. E. & Kirik, O. V. Brain Microglia and Microglial Markers. Neurosci. Behav. Physiol. 46, 284–290 (2016).

45. Ruan, C. & Elyaman, W. A New Understanding of TMEM119 as a Marker of Microglia. Front. Cell. Neurosci. 16, (2022).

46. Broussard, J. I. et al. Repeated mild traumatic brain injury produces neuroinflammation, anxiety-like behaviour and impaired spatial memory in mice. Brain Inj. 32, 113–122 (2018).

47. Porsolt, R. D., Brossard, G., Hautbois, C. & Roux, S. Rodent Models of Depression: Forced Swimming and Tail Suspension Behavioral Despair Tests in Rats and Mice. Curr. Protoc. Neurosci. 14, 8.10A.1–8.10A.10 (2001).

48. Steru, L., Chermat, R., Thierry, B. & Simon, P. The tail suspension test: A new method for screening antidepressants in mice. Psychopharmacology (Berl.) 85, 367–370 (1985).

49. Cryan, J. F., Mombereau, C. & Vassout, A. The tail suspension test as a model for assessing antidepressant activity: review of pharmacological and genetic studies in mice. Neurosci. Biobehav. Rev. 29, 571–625 (2005).

50. Lab, B. BuckleyLabEmory/TSTScoreHelper. (2025).

51. Padilla, E. et al. Novelty-evoked activity in open field predicts susceptibility to helpless behavior. Physiol. Behav. 101, 746–754 (2010).

52. McGowan, N. M. & Coogan, A. N. Circadian and behavioural responses to shift work-like schedules of light/dark in the mouse. J. Mol. Psychiatry 1, 7 (2013).

53. Lueptow, L. M. Novel Object Recognition Test for the Investigation of Learning and Memory in Mice. J. Vis. Exp. JoVE 55718 (2017) doi:10.3791/55718.

54. Lee, S. Y., Zheng, C., Brothers, R. & Buckley, E. M. Small separation frequency-domain near-infrared spectroscopy for the recovery of tissue optical properties at millimeter depths. Biomed. Opt. Express 10, 5362–5377 (2019).

55. Coley, A. A. & Gao, W.-J. PSD-95 deficiency disrupts PFC-associated function and behavior during neurodevelopment. Sci. Rep. 9, 9486 (2019).

56. Béïque, J.-C. & Andrade, R. PSD-95 regulates synaptic transmission and plasticity in rat cerebral cortex. J. Physiol. 546, 859–867 (2003).

57. Kettenmann, H., Hanisch, U.-K., Noda, M. & Verkhratsky, A. Physiology of microglia. Physiol. Rev. 91, 461–553 (2011).

58. Buckley, E. M. Decreased microvascular cerebral blood flow assessed by diffuse correlation spectroscopy after repetitive concussions in mice. J Cereb Blood Flow Metab J Int Soc Cereb Blood Flow Metab 35, 1995–2000 (2015).

59. Sankar, S. B. Low cerebral blood flow is a non-invasive biomarker of neuroinflammation after repetitive mild traumatic brain injury. Neurobiol Dis 124, 544–554 (2019).

60. Baskin, B. M. et al. Timing matters: Sex differences in inflammatory and behavioral outcomes following repetitive blast mild traumatic brain injury. Brain. Behav. Immun. 110, 222–236 (2023).

61. Tucker, L. B., Velosky, A. G., Fu, A. H. & McCabe, J. T. Chronic Neurobehavioral Sex Differences in a Murine Model of Repetitive Concussive Brain Injury. Front. Neurol. 10, (2019).

62. Tucker, L. B. et al. Sex differences in cued fear responses and parvalbumin cell density in the hippocampus following repetitive concussive brain injuries in C57BL/6J mice. PLOS ONE 14, e0222153 (2019).

63. Bazarian, J. J., Blyth, B., Mookerjee, S., He, H. & McDermott, M. P. Sex Differences in Outcome after Mild Traumatic Brain Injury. J. Neurotrauma 27, 527–539 (2010).

64. Mikolic, A. et al. Explaining Outcome Differences between Men and Women following Mild Traumatic Brain Injury. J. Neurotrauma 38, 3315–3331 (2021).

65. Covassin, T., Elbin, R. J., Bleecker, A., Lipchik, A. & Kontos, A. P. Are there differences in neurocognitive function and symptoms between male and female soccer players after concussions? Am. J. Sports Med. 41, 2890–2895 (2013).

66. Hülskötter, K., Lühder, F., Flügel, A., Herder, V. & Baumgärtner, W. Tamoxifen Application Is Associated with Transiently Increased Loss of Hippocampal Neurons following Virus Infection. Int. J. Mol. Sci. 22, 8486 (2021).

67. Crisci, I. et al. Tamoxifen exerts direct and microglia-mediated effects preventing neuroinflammatory changes in the adult mouse hippocampal neurogenic niche. Glia 72, 1273–1289 (2024).

68. Blum, K. M. et al. Sex and Tamoxifen confound murine experimental studies in cardiovascular tissue engineering. Sci. Rep. 11, 8037 (2021).

69. Álvarez-Aznar, A. et al. Tamoxifen-independent recombination of reporter genes limits lineage tracing and mosaic analysis using CreERT2 lines. Transgenic Res. 29, 53–68 (2020).

70. Yang, H. et al. SIRT1 plays a neuroprotective role in traumatic brain injury in rats via inhibiting the p38 MAPK pathway. Acta Pharmacol. Sin. 38, 168–181 (2017).

71. Bruchas, M. R. et al. Selective p38α MAPK deletion in serotonergic neurons produces stress-resilience in models of depression and addiction. Neuron 71, 498–511 (2011).

72. Stefanoska, K. et al. Neuronal MAP kinase p38α inhibits c-Jun N-terminal kinase to modulate anxiety-related behaviour. Sci. Rep. 8, 14296 (2018).

73. Whitfield, D. R. et al. Assessment of ZnT3 and PSD95 protein levels in Lewy body dementias and Alzheimer’s disease: association with cognitive impairment. Neurobiol. Aging 35, 2836–2844 (2014).

74. McEachern, E. P., Coley, A. A., Yang, S.-S. & Gao, W.-J. PSD-95 deficiency alters GABAergic inhibition in the prefrontal cortex. Neuropharmacology 179, 108277 (2020).

75. Bustos, F. J. et al. PSD95 Suppresses Dendritic Arbor Development in Mature Hippocampal Neurons by Occluding the Clustering of NR2B-NMDA Receptors. PLOS ONE 9, e94037 (2014).

76. Alam, J., Blackburn, K. & Patrick, D. Neflamapimod: Clinical Phase 2b-Ready Oral Small Molecule Inhibitor of p38α to Reverse Synaptic Dysfunction in Early Alzheimer’s Disease. J. Prev. Alzheimers Dis. 4, 273–278 (2017).

77. Falcicchia, C., Tozzi, F., Arancio, O., Watterson, D. M. & Origlia, N. Involvement of p38 MAPK in Synaptic Function and Dysfunction. Int. J. Mol. Sci. 21, 5624 (2020).

78. Larson, K., Damon, M., Randhi, R., Nixon-Lee, N. & Dixon, K. J. Selective Inhibition of Soluble TNF using XPro1595 Improves Hippocampal Pathology to Promote Improved Neurological Recovery Following Traumatic Brain Injury in Mice. http://www.eurekaselect.com https://www.eurekaselect.com/article/124336.

79. Khuman, J. Tumor necrosis factor alpha and Fas receptor contribute to cognitive deficits independent of cell death after concussive traumatic brain injury in mice. J Cereb Blood Flow Metab J Int Soc Cereb Blood Flow Metab 31, 778–789 (2011).

80. Newell, E. A. et al. Combined Blockade of Interleukin-1α and −1β Signaling Protects Mice from Cognitive Dysfunction after Traumatic Brain Injury. eNeuro 5, ENEURO.0385-17.2018 (2018).

81. Flygt, J. et al. Neutralization of Interleukin-1β following Diffuse Traumatic Brain Injury in the Mouse Attenuates the Loss of Mature Oligodendrocytes. J. Neurotrauma 35, 2837–2849 (2018).

82. Dash, P. K. et al. Activation of Alpha 7 Cholinergic Nicotinic Receptors Reduce Blood–Brain Barrier Permeability following Experimental Traumatic Brain Injury. J. Neurosci. 36, 2809–2818 (2016).

83. Ho, M.-H. et al. CCL5 via GPX1 activation protects hippocampal memory function after mild traumatic brain injury. Redox Biol. 46, 102067 (2021).

84. Ho, M.-H. et al. CCL5/RANTES signaling in inflammation dysregulation after mild traumatic brain injury. J. Biomed. Sci. 33, 10 (2026).

85. Cherry, J. D. et al. CCL11 is increased in the CNS in chronic traumatic encephalopathy but not in Alzheimer’s disease. PLOS ONE 12, e0185541 (2017).

86. Gao, W. et al. IL-2/Anti-IL-2 Complex Attenuates Inflammation and BBB Disruption in Mice Subjected to Traumatic Brain Injury. Front. Neurol. 8, (2017).

87. Johnson, E. A. et al. Increased expression of the chemokines CXCL1 and MIP-1α by resident brain cells precedes neutrophil infiltration in the brain following prolonged soman-induced status epilepticus in rats. J. Neuroinflammation 8, 41 (2011).

88. Zhu, X. et al. Upregulation of CCL3/MIP-1alpha regulated by MAPKs and NF-kappaB mediates microglial inflammatory response in LPS-induced brain injury. Acta Neurobiol. Exp. (Warsz.) 76, 304–317 (2016).

89. Hall, C. N. et al. Capillary pericytes regulate cerebral blood flow in health and disease. Nature 508, 55–60 (2014).

90. Lynch, C. M. et al. Nox2-Derived Superoxide Contributes to Cerebral Vascular Dysfunction in Diet-Induced Obesity. Stroke 44, 3195–3201 (2013).

91. Tong, X.-K., Nicolakakis, N., Kocharyan, A. & Hamel, E. Vascular Remodeling versus Amyloid β-Induced Oxidative Stress in the Cerebrovascular Dysfunctions Associated with Alzheimer’s Disease. J. Neurosci. 25, 11165–11174 (2005).

92. Wu, R. et al. c-Abl–p38α signaling plays an important role in MPTP-induced neuronal death. Cell Death Differ. 23, 542–552 (2016).

93. Ghatan, S. et al. p38 Map Kinase Mediates Bax Translocation in Nitric Oxide–Induced Apoptosis in Neurons. J. Cell Biol. 150, 335–348 (2000).

94. Pandey, A. K. et al. Neflamapimod induces vasodilation in resistance mesenteric arteries by inhibiting p38 MAPKα and downstream Hsp27 phosphorylation. Sci. Rep. 12, 4905 (2022).

95. Sanchez, A. et al. p38 MAPK: a mediator of hypoxia-induced cerebrovascular inflammation. J. Alzheimers Dis. JAD 32, 587–597 (2012).

96. Ni, Y. et al. TNFα alters occludin and cerebral endothelial permeability: Role of p38MAPK. PLoS ONE 12, e0170346 (2017).

97. Armstead, W. M. PTK, ERK and p38 MAPK contribute to impaired NMDA-induced vasodilation after brain injury. Eur. J. Pharmacol. 474, 249–254 (2003).

98. Pan, Y.-X. et al. Intracisternal administration of SB203580, a p38 mitogen-activated protein kinase inhibitor, attenuates cerebral vasospasm via inhibition of tumor-necrosis factor-α. J. Clin. Neurosci. 20, 726–730 (2013).

99. Prendergast, B. J., Onishi, K. G. & Zucker, I. Female mice liberated for inclusion in neuroscience and biomedical research. Neurosci. Biobehav. Rev. 40, 1–5 (2014).

100. Wagner, A. K. et al. Gender associations with chronic methylphenidate treatment and behavioral performance following experimental traumatic brain injury. Behav. Brain Res. 181, 200–209 (2007).

101. Adams, C. et al. Neurogliovascular dysfunction in a model of repeated traumatic brain injury. Theranostics 8, 4824–4836 (2018).

102. Lynch, C. E. et al. Impairment of cerebrovascular reactivity in response to hypercapnic challenge in a mouse model of repetitive mild traumatic brain injury. J. Cereb. Blood Flow Metab. Off. J. Int. Soc. Cereb. Blood Flow Metab. 41, 1362–1378 (2021).

103. Lynch, C. E. et al. Chronic cerebrovascular abnormalities in a mouse model of repetitive mild traumatic brain injury. Brain Inj. 30, 1414–1427 (2016).

