## Supplementary Material for "Neuronal p38a knockout protects against neurological consequences following repetitive mild traumatic brain injury"

### Supplementary Figures

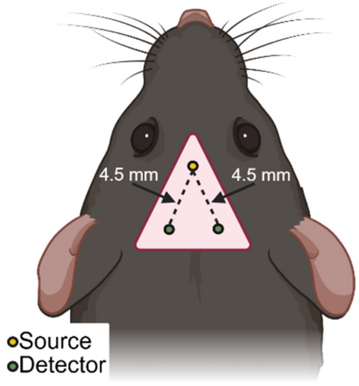


**Fig. S1. Illustration of the diffuse correlation spectroscopy (DCS) sensor geometry and typical placement for measurements**. The DCS sensor (pink triangle) consisted of a single source optode (yellow) spaced equidistance away from 2 detector optodes designed to interrogate regional hemodynamics in the right and left hemisphere. Created in BioRender. Triplett, S. (2026) https://BioRender.com/vtmdajx.

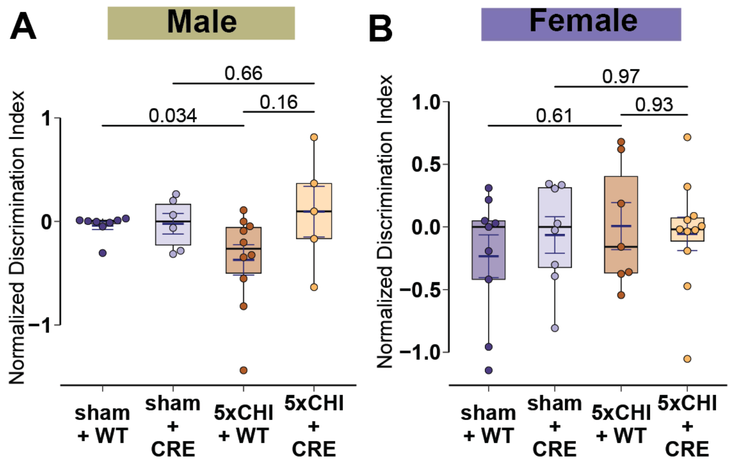


**Fig S2. Neuronal p38α knockout protected against injury-induced short-term memory loss at 4 weeks post injury.** Box plots of the discrimination index on the novel object recognition task in **A.** males (n=5-10/group) and **B**. females (n=7-12/group). Each dot denotes one animal. mean±SEM are also displayed, p-values are estimated from paired Wilcoxon rank sum tests. 5xCHI = 5 once-daily closed-head injuries; WT = p38α$^{fl/fl}$; wildtype control mice; CRE = p38α$^{fl/fl}$; Cre mice.

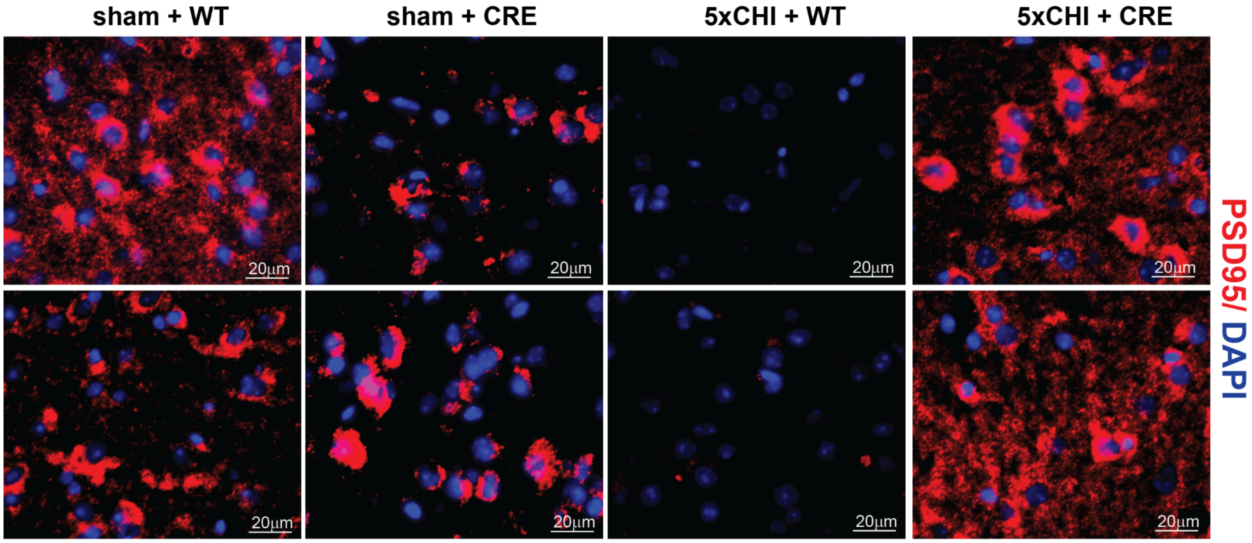


**Fig S3. Immunohistochemistry (IHC) images of PSD95 in frontal cortex at 4 weeks post rmTBI. A**. Additional representative IHC images in frontal cortex showing PSD95 stain (red) and DAPI (blue) (scale bar: 20µm, representative sections from n=3-4 mice/group). IHC = immunohistochemistry. PSD95 = post synaptic marker 95; 5xCHI = 5 once-daily closed-head injuries; WT = $p38\alpha^{fl/fl}$; wildtype control mice; CRE = $p38\alpha^{fl/fl}$; Cre mice.

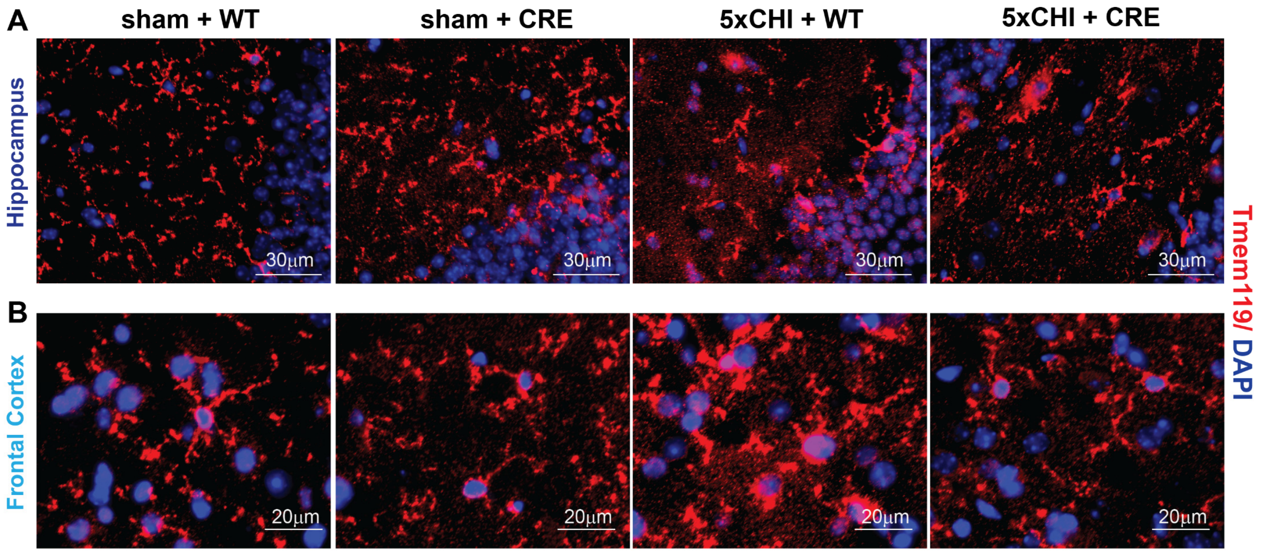


**Fig S4. Immunohistochemistry (IHC) images of Tmem119 at 4 hours post rmTBI. A**. Additional representative IHC images in hippocampus and **B**. frontal cortex showing Tmem119 stain (red) and DAPI (blue) (scale bar: 20µm, representative sections from n=3-4 mice/group). 5xCHI = 5 once-daily closed-head injuries; WT = p38α$^{fl/fl}$; wildtype control mice; CRE = p38α$^{fl/fl}$; Cre mice; Tmem119 = transmembrane protein 119.

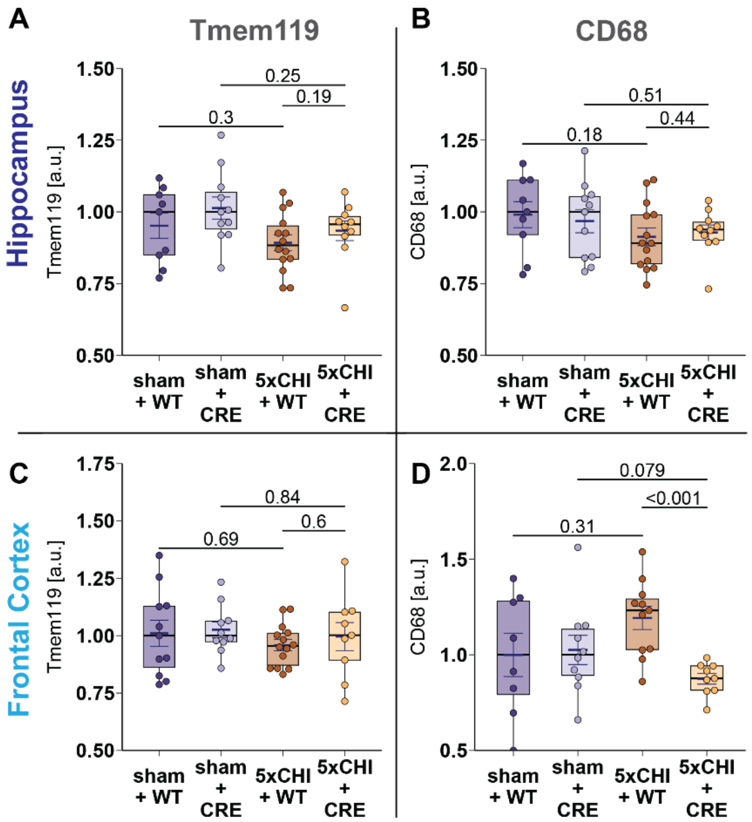


**Fig S5. Effects of 5xCHI and neuronal p38α knockout on microglial reactivity at 4 weeks post rmTBI in males.** The top panels (A, B) represent data from hippocampus and bottom panels (C, D) represent data from frontal cortex (n=9-14/group). **A**. Box plots of ELISA measured (A) hippocampal Tmem119, **B**. hippocampal CD68, **C**. cortical Tmem119, **D**. cortical CD68. All values are normalized to the median of the corresponding sham-injured group for each genotype. Each dot denotes one animal. mean±SEM are also displayed, p-values are estimated from Wilcoxon rank sum tests with Bonferroni adjustment for multiple comparisons. 5xCHI = 5 once-daily closed-head injuries; WT = p38α$^{fl/fl}$; wildtype control mice; CRE = p38α$^{fl/fl}$; Cre mice; ELISA = enzyme-linked immunosorbent assay; Cluster of Differentiation 68; Tmem119 = transmembrane protein 119.

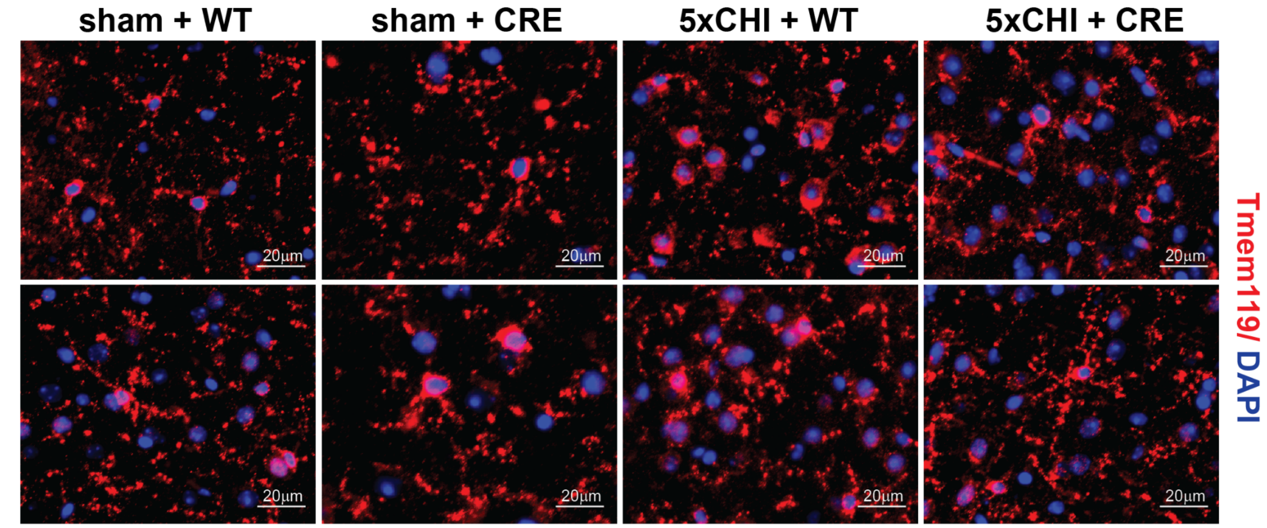


**Fig S6. Immunohistochemistry (IHC) for Tmem119 in frontal cortex at 4 weeks post rmTBI in males. A**. Representative IHC images in frontal cortex showing Tmem119 stain (red) and DAPI (blue) (scale bar: 20µm, representative sections from n=3-4 mice/group). 5xCHI = 5 once-daily closed-head injuries; WT = $p38\alpha^{fl/fl}$; wildtype control mice; CRE = $p38\alpha^{fl/fl}$; Cre mice; IHC = immunohistochemistry; Tmem119 = transmembrane protein 119.

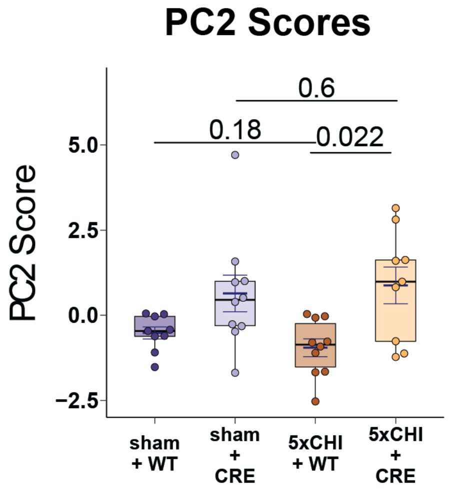


**Fig S7. Principal component analysis of cytokine profiles at 4 hours post rmTBI.** Boxplot of the scores on principal component 2 (PC2) by group. Each dot denotes one animal. mean±SEM are also displayed, p-values are estimated from Wilcoxon rank sum tests with Bonferroni adjustment for multiple comparisons. a.u.= arbitrary units. 5xCHI = 5 once-daily closed-head injuries; WT = p38α$^{fl/fl}$; wildtype control mice; CRE = p38α$^{fl/fl}$; Cre mice.

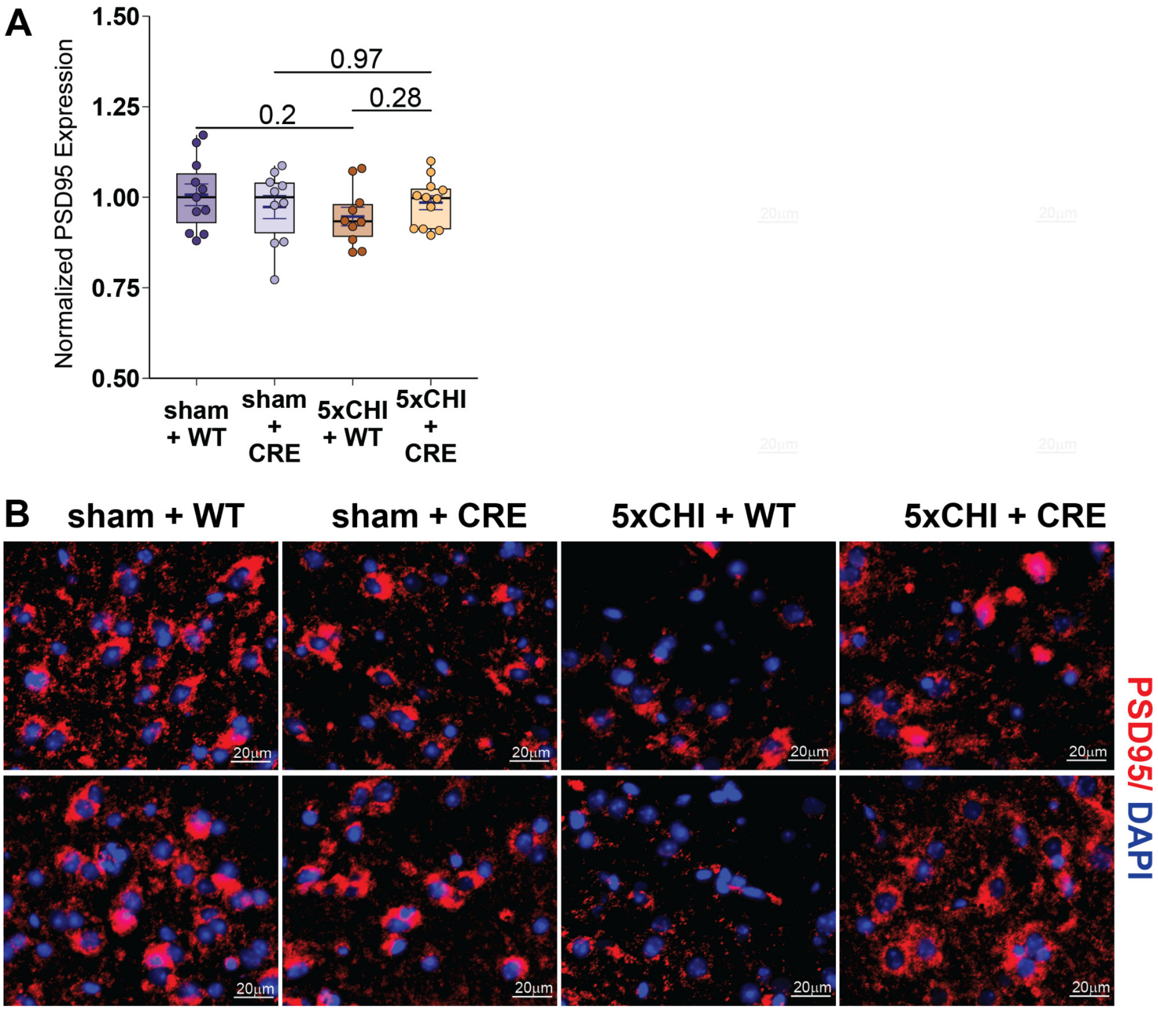


**Fig S8. Effects of 5xCHI and neuronal p38α knockout on synaptic density at 4 weeks post rmTBI in females. A**. Box plot of PSD95 ELISA results (n=10-12 /group). All values reported were normalized by the means of the corresponding sham-injured group for each genotype. Each dot denotes an individual animal. mean±SEM are also displayed, p-values are estimated from Wilcoxon rank sum tests with Bonferroni adjustment for multiple comparisons. **B**. Representative IHC images in frontal cortex showing PSD95 stain (red) and DAPI (blue) (scale bar: 20μm, representative sections from n=3-4 mice/group). a.u.= arbitrary units. IHC = immunohistochemistry. PSD95 = post synaptic marker 95; 5xCHI = 5 once-daily closed-head injuries; WT = p38α$^{fl/fl}$; wildtype control mice; CRE = p38α$^{fl/fl}$; Cre mice.

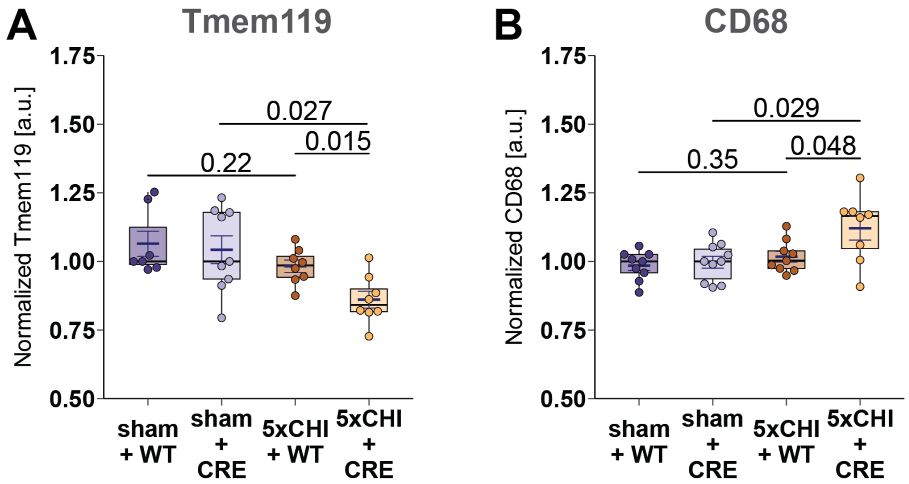


**Fig S9. Effects of 5xCHI and neuronal p38α knockout on microglial reactivity at 4 hours post rmTBI in females.** Boxplots of ELISA-measured **A**. Tmem119 (n=8-10/group) and **B**. CD68 (n=8-10/group). All values are normalized by the median of the corresponding sham-injured group for each genotype. Each dot denotes one animal. mean±SEM are also displayed, p-values are estimated from Wilcoxon rank sum tests with Bonferroni adjustment for multiple comparisons. 5xCHI = 5 once-daily closed-head injuries; WT = p38α$^{fl/fl}$; wildtype control mice; CRE = p38α$^{fl/fl}$; Cre mice; ELISA = enzyme-linked immunosorbent assay; Cluster of Differentiation 68, Tmem119 = transmembrane protein 119.

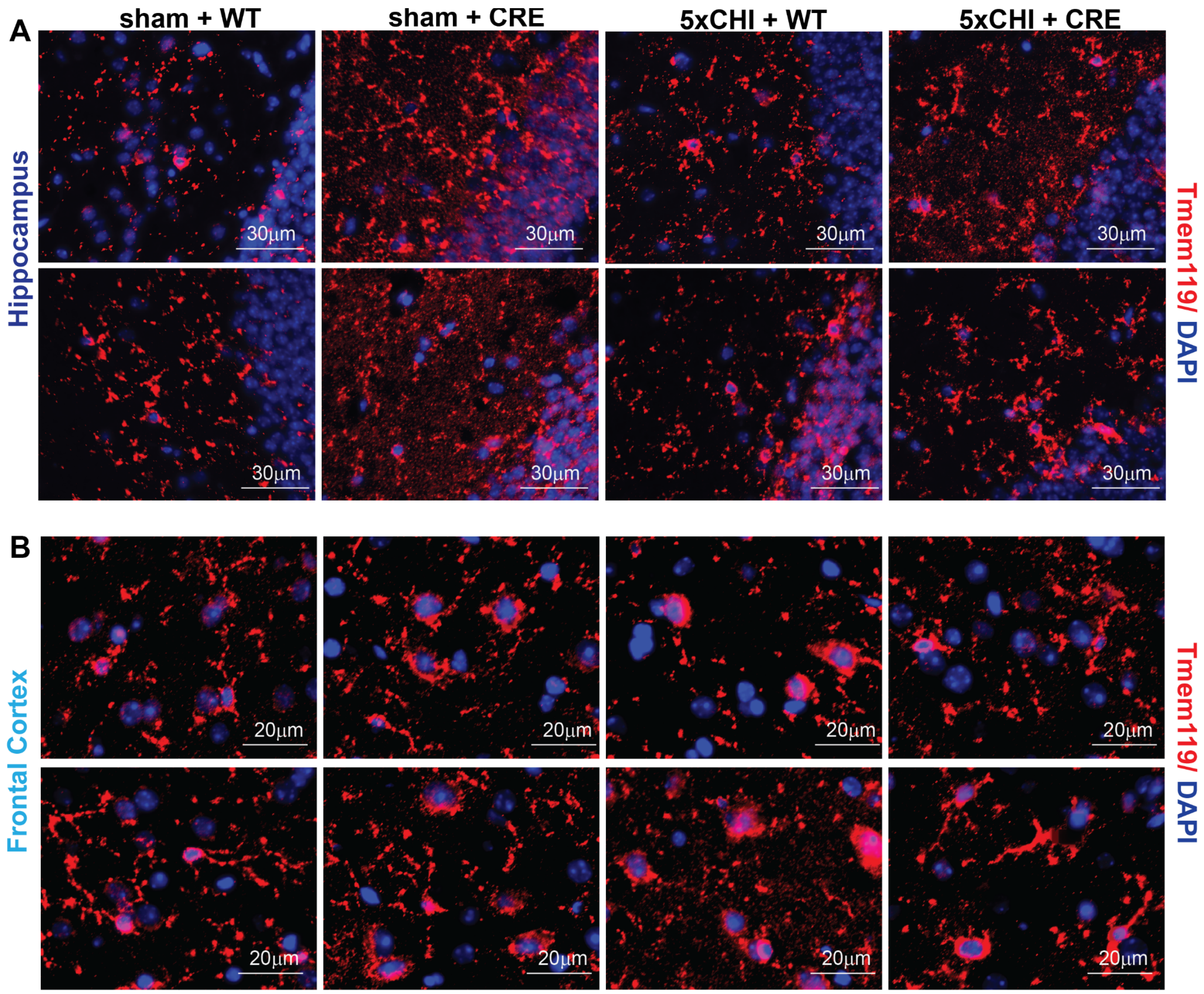


**Fig S10. Immunohistochemistry (IHC) for Tmem119 at 4 hours post rmTBI in females. A**. Representative IHC images in hippocampus and **B**. frontal cortex showing Tmem119 stain (red) and DAPI (blue) (scale bar: 30µm, representative sections from n=3-4 mice/group). 5xCHI = 5 once-daily closed-head injuries; WT = p38α$^{fl/fl}$; wildtype control mice; CRE = p38α$^{fl/fl}$; Cre mice; Tmem119 = transmembrane protein 119.

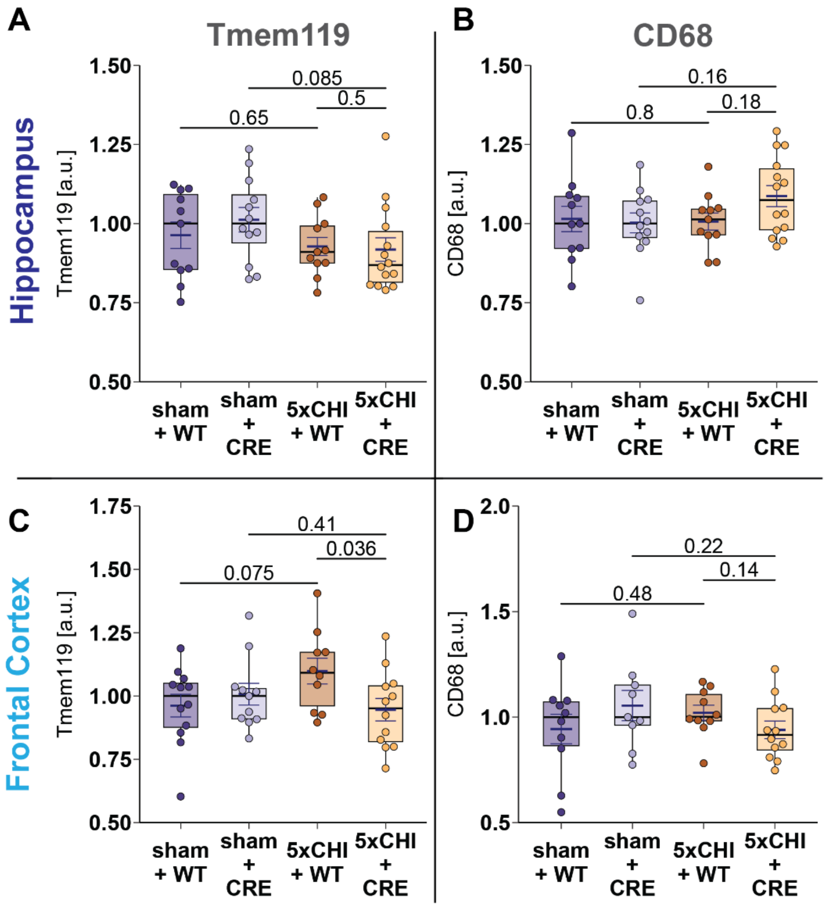


**Fig S11. Effects of 5xCHI and neuronal p38α knockout on microglial reactivity at 4 weeks post rmTBI in females.** The top panels (A, B) represent data from hippocampus and bottom panels (C, D) represent data from frontal cortex (n=9-14/group). **A**. Box plots of ELISA measured (A) hippocampal Tmem119, **B**. hippocampal CD68, **C**. cortical Tmem119, **D**. cortical CD68. All values reported were normalized by the median of the corresponding sham-injured group for each genotype. Each dot denotes one animal. mean±SEM are also displayed, p-values are estimated from Wilcoxon rank sum tests with Bonferroni adjustment for multiple comparisons. 5xCHI = 5 once-daily closed-head injuries; WT = p38α$^{fl/fl}$; wildtype control mice; CRE = p38α$^{fl/fl}$; Cre mice. ELISA = enzyme-linked immunosorbent assay, Cluster of Differentiation 68, Tmem119 = transmembrane protein 119.

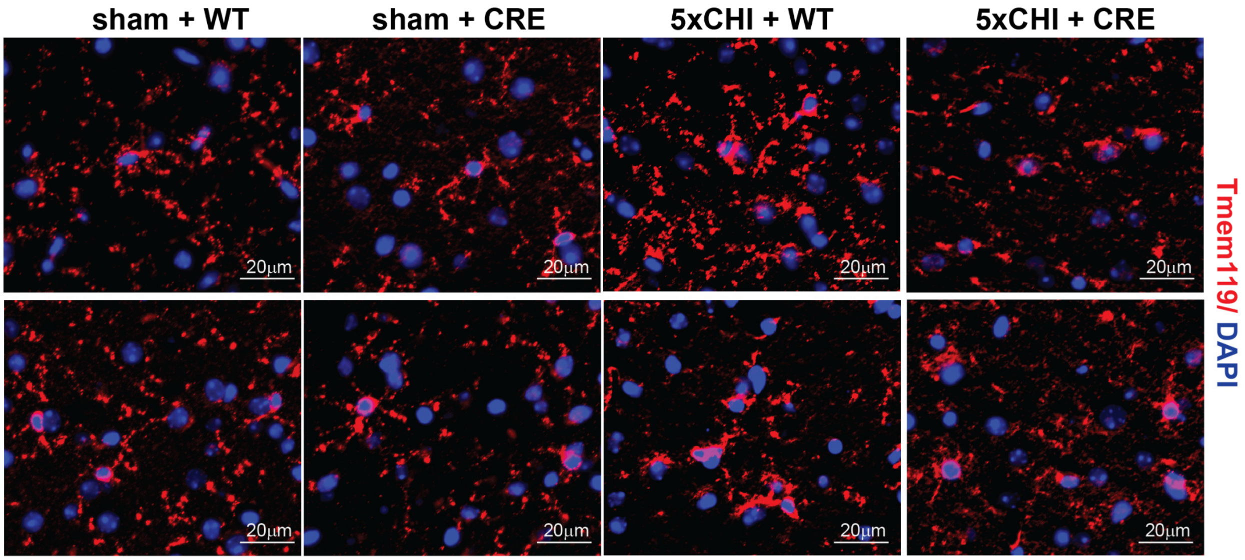


**Fig S12. Immunohistochemistry (IHC) for Tmem119 in frontal cortex at 4 weeks post rmTBI in females. A**. Representative IHC images in cortex showing Tmem119 stain (red) and DAPI (blue) (scale bar: 20μm, representative sections from n=3-4 mice/group). 5xCHI = 5 once-daily closed-head injuries; WT = p38α$^{fl/fl}$; wildtype control mice; CRE = p38α$^{fl/fl}$; Cre mice; Tmem119 = transmembrane protein 119.

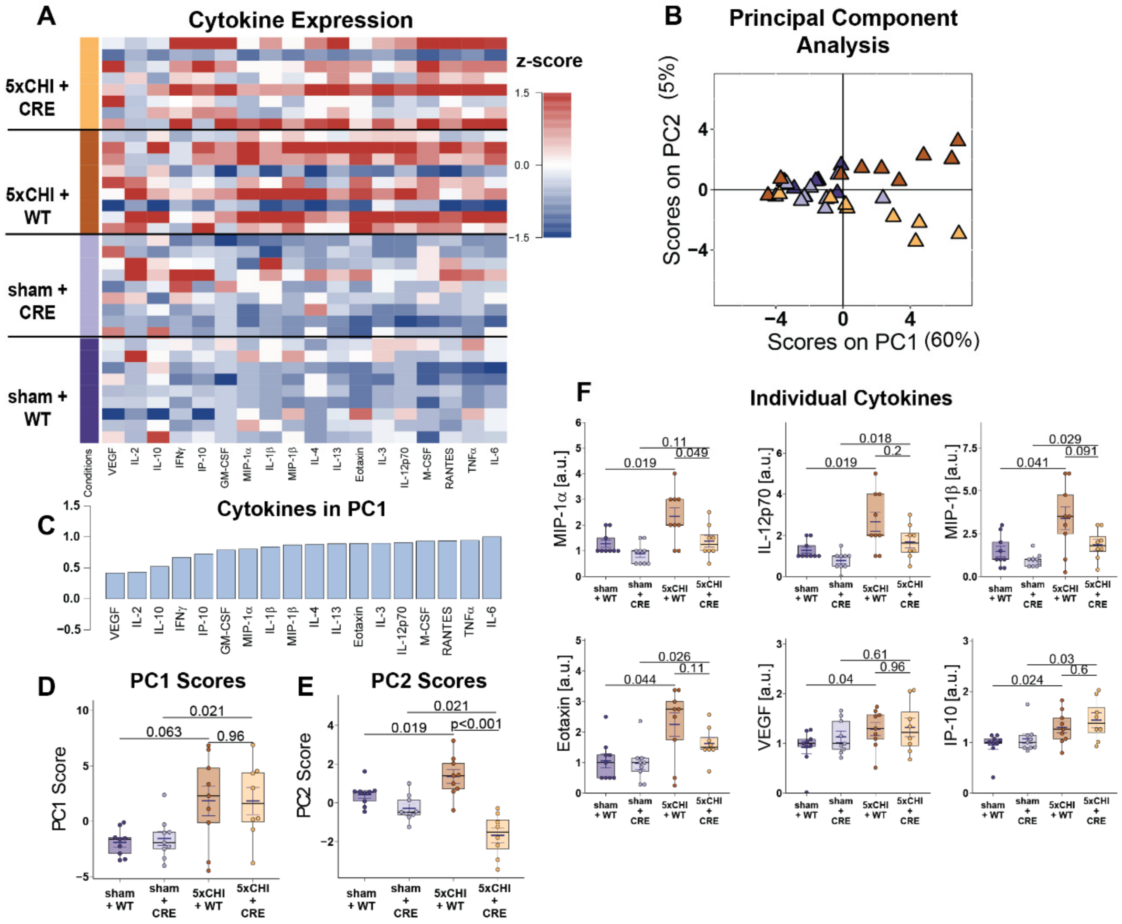


**Fig S13. Neuronal p38α knockout protected against injury-induced elevation in cytokine expression at 4 hours post rmTBI in females. A**. Luminex analysis of 18 hippocampal cytokines at 4 hours after 5xCHI or sham-injury (n=9-10). Each column represents one cytokine analyte, while each row denotes an individual animal (columns are z-scored) **B**. Scores on principal component 1 (PC1) seperated the 5xCHI+WT group to the right. Each triangle denotes one animal. **C**. Boxplot of the scores on PC1 by group (Wilcoxon rank sum tests). **D**. PC1 consisted of a profile of cytokines that were positively correlated with PC1 and thus the 5xCHI+WT group in **C**. **E**. Boxplot of the scores on PC2 by group. **F**. Box plots for individual cytokines. All values reported were normalized by the median of the corresponding sham-injured group for each genotype. Each dot denotes one animal. mean±SEM are also displayed, p-values are estimated from Wilcoxon rank sum tests with Bonferroni adjustment for multiple comparisons. a.u.= arbitrary units. 5xCHI = 5 once-daily closed-head injuries; WT = p38α$^{fl/fl}$; wildtype control mice; CRE = p38α$^{fl/fl}$; Cre mice.

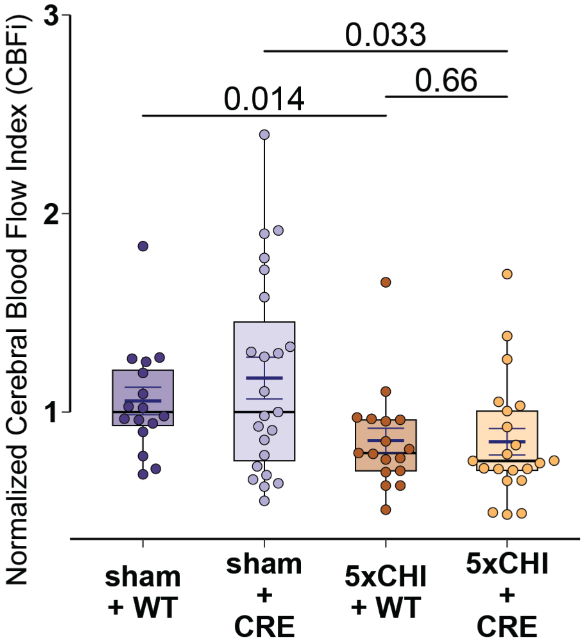


**Fig S14. Neuronal p38α knockout had limited effect on injury-induced decreases in cerebral blood flow (CBF) in females. A.** Box plot of the normalized cerebral blood flow index (CBFi) at 4 hours after 5xCHI or sham-injury dichotomized by group (n=16-23/group). All values were normalized by the median of the corresponding sham-injured group for each genotype. Each dot denotes one animal. mean±SEM are also displayed, p-values are estimated from Wilcoxon rank sum tests with Bonferroni adjustment for multiple comparisons. 5xCHI = 5 once-daily closed-head injuries; WT = p38αfl/fl; wildtype control mice; CRE = p38αfl/fl; Cre mice.

### Supplementary Tables

| Analyte | **sham + WT vs. 5xCHI + WT** | **sham + CRE vs. 5xCHI + CRE** | **5xCHI + WT vs. 5xCHI + CRE** |
|---|---|---|---|
| Eotaxin | 0.004 | ns | <0.001 |
| GM-CSF | 0.012 | ns | ns |
| IFNγ | ns | ns | ns |
| IL-10 | ns | ns | ns |
| IL-12p70 | 0.005 | ns | ns |
| IL-13 | 0.013 | ns | 0.03 |
| IL-1β | ns | ns | 0.028 |
| IL-2 | <0.001 | ns | 0.01 |
| IL-3 | 0.016 | ns | 0.043 |
| IL-4 | ns | ns | 0.014 |
| IL-6 | 0.011 | ns | ns |
| IP-10 | 0.007 | ns | 0.03 |
| M-CSF | 0.004 | ns | 0.018 |
| MIP-1α | 0.015 | ns | <0.001 |
| MIP-1β | 0.012 | ns | ns |
| RANTES | 0.016 | ns | 0.01 |
| TNFα | 0.007 | ns | 0.033 |
| VEGF | ns | ns | ns |

**Table S1. p-values for male cytokine expression differences by analyte.** All p-values are estimated from Wilcoxon rank sum tests with Bonferroni adjustment for multiple comparisons. ns = not significant; 5xCHI = 5 once-daily closed-head injuries; WT = $p38\alpha^{fl/fl}$; wildtype control mice; CRE = $p38\alpha^{fl/fl}$; Cre mice.

| Analyte | **sham + WT vs. 5xCHI + WT** | **sham + CRE vs. 5xCHI + CRE** | **5xCHI + WT vs. 5xCHI + CRE** |
|---|---|---|---|
| Eotaxin | 0.044 | 0.026 | ns |
| GM-CSF | ns | 0.023 | ns |
| IFNγ | ns | ns | 0.008 |
| IL-10 | ns | ns | ns |
| IL-12p70 | 0.019 | 0.018 | ns |
| IL-13 | ns | 0.014 | ns |
| IL-1β | ns | ns | ns |
| IL-2 | ns | ns | ns |
| IL-3 | 0.035 | ns | ns |
| IL-4 | ns | ns | ns |
| IL-6 | ns | 0.047 | ns |
| IP-10 | 0.024 | 0.03 | ns |
| M-CSF | 0.007 | ns | ns |
| MIP-1α | 0.019 | 0.11 | 0.049 |
| MIP-1β | 0.041 | 0.029 | ns |
| RANTES | ns | ns | ns |
| TNFα | ns | 0.027 | ns |
| VEGF | 0.04 | ns | ns |

**Table S2. p-values for female cytokine expression differences by analyte.** All p-values are estimated from Wilcoxon rank sum tests with Bonferroni adjustment for multiple comparisons. ns = not significant; 5xCHI = 5 once-daily closed-head injuries; WT = p38α$^{fl/fl}$; wildtype control mice; CRE = p38α$^{fl/fl}$; Cre mice.
